# CRISPR-Cas9 Induced Knockout of *BEL5* in Tetraploid Potato: Optimized Methodology via Repeated *de novo* Regeneration and Impact on Tuberization

**DOI:** 10.64898/2026.07.21.739726

**Authors:** Andrea Zounková, Daniele Chirivì, Adéla Přibylová, Damiano Martignago, Jitka Myslivcová, Tomáš Mašek, Lukáš Fischer, Camilla Betti, Fabio Fornara, Petra Mašková

## Abstract

CRISPR-Cas9 has emerged as a powerful tool for targeted genome editing in plants; however, its application in tetraploid potato (*Solanum tuberosum* ssp. *tuberosum*) remains challenging due to its vegetative propagation and complex highly heterozygous genome. Availability of whole-genome sequence data for the specific genotype is crucial to ensure complete knockout of all alleles of target genes while minimizing off-target mutations. In this study, using the tetraploid potato cultivar Désirée, we report, a complete CRISPR-Cas9-mediated knockout of the *BEL5* gene, encoding a transcription factor, known as one of the key regulators driving tuber formation. We employed *Agrobacterium*-mediated transformation and demonstrated that repeated *de novo* regeneration could improve editing efficiency by promoting emergence of new mutations. *BEL5* knockout plants exhibited a delayed onset of tuberization under inductive short-day conditions in hydroponics; however, their overall tuber yields were comparable to wild type plants. Based on our results, we propose a regulatory role of BEL5 in the timing of tuber onset but, unexpectedly, its dispensability for tuber development in modern cultivated potato. Besides providing functional insight into the BEL5 role in potato, this study includes a methodological approach for efficient CRISPR-Cas9 gene editing in this vegetatively propagated polyploid crop, along with strategies for detecting mutations in genes that lack clear phenotypic manifestation.

## 1. Introduction

Potato (*Solanum tuberosum*) is the fourth most important crop worldwide, playing a key role in global food security (Devaux *et al*., 2020). Considering challenges such as climate change and population growth, developing new potato varieties with higher yields, improved disease resistance, and better environmental adaptability is crucial (Del Mar Martínez-Prada *et al*., 2021). However, breeding through conventional methods is slow, laborious and imprecise. Especially, stacking multiple traits within one genotype is hardly achievable without targeted genome editing. CRISPR-Cas9 has revolutionized plant breeding by enabling precise, targeted modifications in the genome. This technology has been widely applied in the model plant *Arabidopsis thaliana*, but its use has also expanded to various crop species (Cardi *et al*., 2023). In potato, CRISPR-Cas9 methodology started to be adopted particularly for inducing targeted gene knockouts (Tiwari *et al*., 2022). However, achieving complete gene knockout remains a significant challenge, as only a few published studies reliably demonstrate successful gene knockout in cultivated potato, with typically low efficiency e.g. (Kieu *et al*., 2021; Lebedeva *et al*., 2022; Norouzi *et al*., 2024; Sevestre *et al*., 2020; Takeuchi *et al*., 2021). Cultivated potato (*S. tuberosum* ssp. *tuberosum*) possesses a highly heterozygous autotetraploid genome, making targeted gene editing particularly challenging. To address these challenges, many studies rely on diploid potato varieties, such as *S. tuberosum* group *phureja*, or on homozygous and heterozygous doubled haploids, to increase the probability of obtaining fully edited lines and simplify genetic analysis (Eggers *et al*., 2021; Enciso-Rodriguez *et al*., 2019; Tang *et al*., 2022; Wang *et al*., 2015; Ye *et al*., 2018). However, these synthetic genotypes or landraces are genetically distant from the widely cultivated tetraploid genotypes, limiting the applicability of the insights gained into breeding settings (Gutaker *et al*., 2019; Hardigan *et al*., 2017).

A prerequisite for effective CRISPR-Cas9-based gene editing in potato is the design of gRNAs that specifically target all the alleles but also minimize the risk of off-target mutations. This requires access to whole-genome sequencing data for the specific cultivar, as potatoes exhibit one of the highest genomic diversities among sequenced crops (Hardigan *et al*., 2017). There are short paired-end Illumina reads data available for a wide range of potato genomes, including those from wild species, landraces, historical herbarium specimens, as well as modern cultivars (Gutaker *et al*., 2019; Hardigan *et al*., 2017; Sevestre *et al*., 2020). While these data offer broad genomic coverage and are especially useful for identifying single nucleotide polymorphisms (SNPs), they have considerable limitations, particularly in accurate allele assignment. The genome-scale data count from the first reference genome of the doubled monoploid potato group *phureja* (clone DM1-3 516 R44, hereafter referred to as DM) till chromosome-scale, haplotype-resolved genomes assemblies of tetraploid potato cultivars (table S1). Haplotype-resolved whole-genome data offer significant advantages in this regard, but they are currently available only for a limited number of genotypes.

Beyond gRNA design, the complexity of potato genome substantially complicates also other steps required to obtain fully edited lines with knockout of all alleles, and this must be carefully considered when selecting appropriate methodologies. Due to high allelic diversity leading to largely unpredictable segregation patterns, together with reduced sexual fertility, potato is usually propagated vegetatively (Bisognin, 2011). Consequently, obtaining homozygous mutants in subsequent generations is generally not feasible, so transformation strategies must aim for editing of all alleles in all cells of the regenerated plant. Protoplast-based editing offers a potential solution; however, plant regeneration from protoplasts is frequently associated with large-scale genomic instability (Fossi *et al*., 2019). Therefore, it is necessary to focus on improving other transformation strategies and approaches for enhancing editing efficiency.

Another challenge arising from the complexity of the potato genome is the reliable identification of induced mutations in all alleles (Sánchez-Gómez *et al*., 2023). This is particularly difficult for genes whose knockouts do not result in a clear phenotypic manifestation. Therefore, a careful selection of methods used for detection of all editing variants across all alleles and/or cells is indispensable for correct interpretation of phenotypic changes.

In this study, we applied CRISPR-Cas9–mediated gene editing to investigate the regulatory pathway controlling potato tuberization. Tuberization is a morphogenetic process characterized by the development of storage organs called tubers from the tips of underground lateral stems, known as stolons (Xu *et al*., 1998). Under inductive conditions, tuber formation is initiated by mobile signals which are produced in leaves, and then transported via phloem to stolons (Bao *et al*., 2025). Among these mobile signals, FLOWERING LOCUS T homologues, particularly SELF-PRUNING 6A (SP6A), and BEL5, a transcription factor, play a central role in positive regulation of tuber initiation (Chen *et al*., 2003; Navarro *et al*., 2011). SP6A is transported as a protein, whereas *BEL5* transcript is stabilised and transported in polypyrimidine tract-binding protein–containing RNPs (Banerjee, *et al*., 2006a; Cho *et al*., 2015; Navarro *et al*., 2011). In stolons, these tuberigenic signals form regulatory complexes: TAC complex (composed of SP6A, FD-like, and 14-3-3 proteins) and BEL5–Knotted1-like homeobox protein (KNOX) complex, which regulate the expression of target genes responsible for tuber development (Sharma *et al*., 2016; Teo *et al*., 2016). Here, we focused on BEL5, whose function has previously been investigated in transgenics with altered *BEL5* expression (Banerjee, *et al*., 2006a,b; Banerjee *et al*., 2009; Chen *et al*., 2003; Cho *et al*., 2015; Kondhare *et al*., 2023; Lin *et al*., 2013; Sharma *et al*., 2016), while a complete knockout of this gene had not yet been reported. Employing stable *Agrobacterium*-mediated transformation, followed by repeated *de novo* regeneration, enhancing editing efficiency, we achieved a complete *BEL5* knockout in the tetraploid potato cultivar Désirée. Phenotypic analysis revealed that inactivation of the *BEL5* gene did not prevent tuberization but delayed tuber onset in mutants. The total tuber yields were comparable to the wild type (WT), suggesting that BEL5 transcriptionally regulates the timing of tuberization onset rather than later stages of tuber development. Moreover, since the methodology for achieving complete gene knockout lines in tetraploid potato is not yet well established, we included a detailed methodological guide for effective CRISPR-Cas9 mediated mutagenesis.

## 2. Results

### 2.1. Optimized step-by-step protocol for CRISPR-Cas9 induced gene knockout in tetraploid potato

In the following sections, we describe the key steps to achieve a complete CRISPR-Cas9–induced gene knockout in cultivated potato (see Fig. 1 for a schematic workflow overview).

**Fig. 1:**
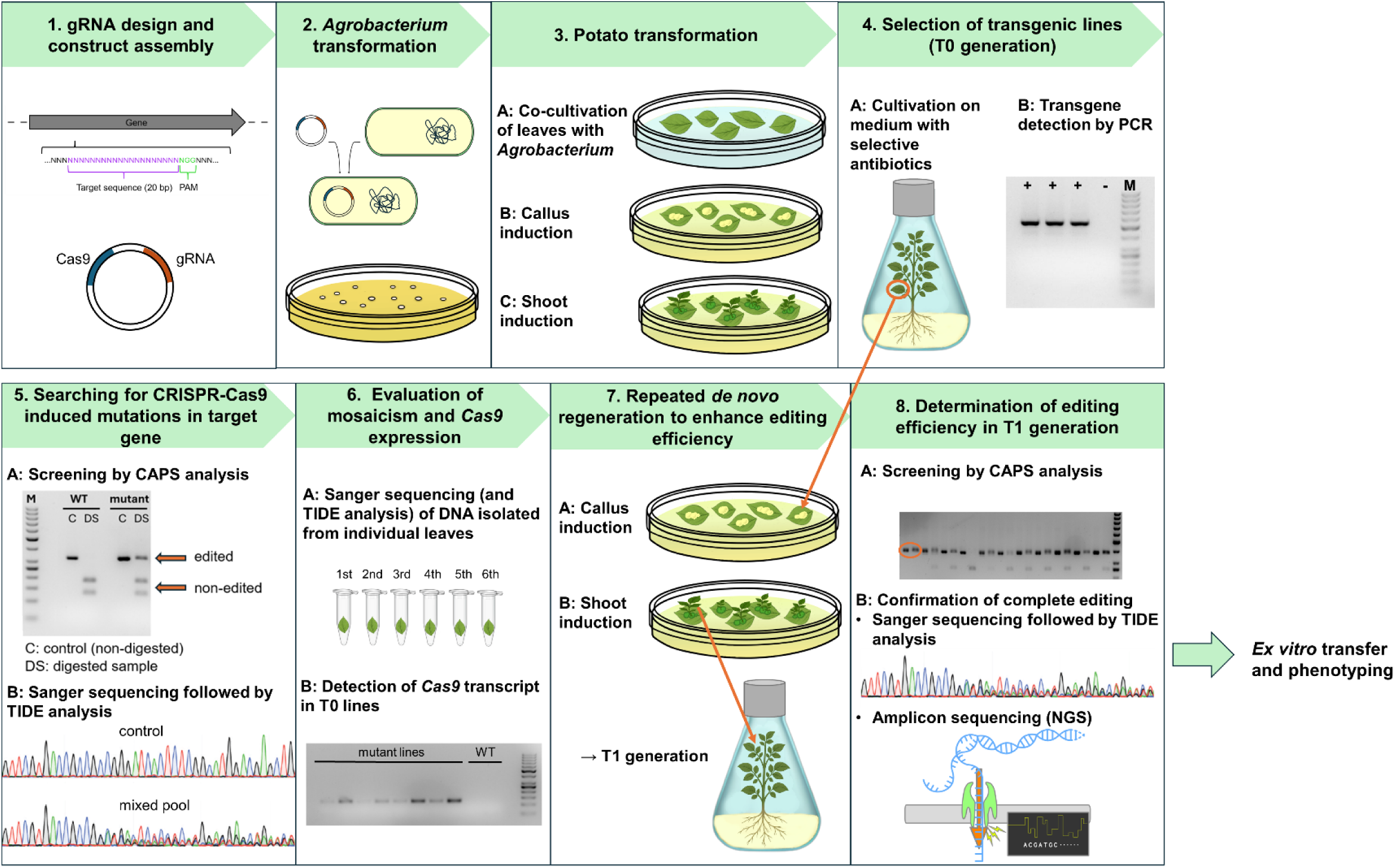
Schematic overview of the workflow for CRISPR-Cas9–induced gene knockout in tetraploid potato, employing repeated *de novo* regeneration to increase editing efficiency: 1) gRNAs targeting all alleles of the gene of interest with minimal off-target risk are designed and the recombinant plasmid assembled; 2) The assembled vector is electroporated into *Agrobacterium tumefaciens*; 3) *Agrobacterium tumefaciens*-mediated stable potato transformation utilizing *de novo* organogenesis; 4) Transgenic lines are selected using cultivation on antibiotic-containing medium, and the presence of the transgene verified by PCR; 5) Edited plant lines are screened for mutations by CAPS analysis using restriction enzymes with recognition sequence in gRNA target sites, and mutation types and their frequency are evaluated by Sanger sequencing with TIDE analysis; 6) Mosaicism and *Cas9* expression are checked in lines selected for repeated *de novo* regeneration; 7) Repeated *de novo* regeneration to promote the introduction of new mutations, thereby increasing editing efficiency; 8) Editing efficiency in the T1 generation determined using CAPS and Sanger sequencing with TIDE analysis, and complete gene knockout in selected T1 lines confirmed by Nanopore sequencing of the full length-gene PCR amplicon, enabling sensitive detection of low-frequency mutations and quantitative and accurate assignment of mutations to individual alleles.

#### 2.1.1. Potato variety selection and identification of target gene alleles

We selected two commercially cultivated tetraploid potato varieties, Désirée and Otava. For Désirée, fully assembled genome sequence was not available at the time of our experiments, so we relied on unassembled Illumina sequencing whole-genome data (Sevestre *et al*., 2020). For Otava, we used already available haplotype-resolved whole-genome data (Sun *et al*., 2022). Since no functional annotation for the target gene *BEL5* has been assigned in either cultivar, we identified four alleles using the *BEL5* sequence from the DM reference genome. In the Otava genome, we used the *BEL5* sequence from the DM reference genome to identify four *BEL5* alleles through BLAST analysis (Table S2). For the cultivar Désirée, we reconstructed the sequences of the four *BEL5* alleles by mapping published Illumina short-reads data onto the *BEL5* sequence from the DM reference genome. Then, we validated the results through amplicon Nanopore sequencing (using Oxford Nanopore Technologies) of PCR products obtained with primers annealing to all *BEL5* alleles at 5’and 3’UTRs (GenBank accession numbers PZ132913-PZ132916 for alleles 1-4, respectively).

#### 2.1.2. Vector selection, gRNA design and transformation

We selected two suitable gRNAs that target the 1st and 4th exon (Fig. 2) of all four *BEL5* alleles in both selected cultivars (Fig. S1). Off-target effects were predicted to be highly unlikely in cv. Otava (Table S3). For Desirée, the prediction of off-target was more difficult as it was based only on unassembled short Illumina reads (Sevestre *et al*., 2020). We did not detect any off-targets with fewer than five mismatches for either gRNA. The only candidate off-target (gRNA 1) was found in transcript XM_006364777.2 (from the reference DM genotype). Given that one mismatch occurs four nucleotides upstream of the PAM and that the extensive mismatching at the 5′ end would hinder cleavage, the likelihood of Cas9 binding with subsequent cutting at this site is minimal (Table S3).

**Fig. 2:**
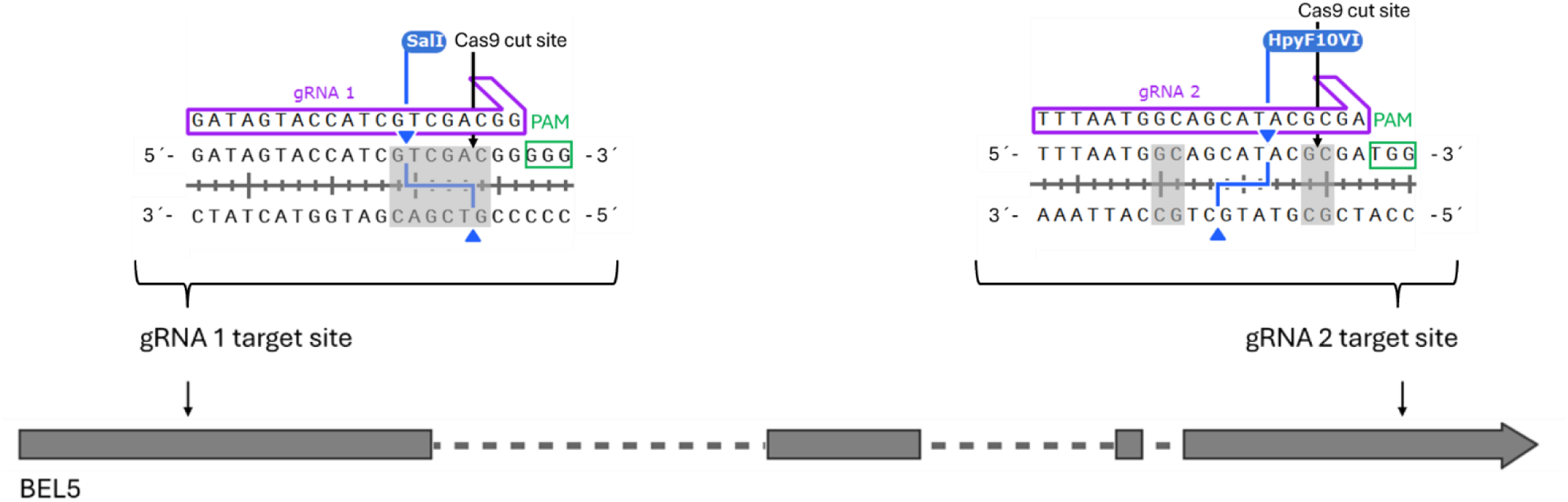
*BEL5* gene-specific gRNAs: gRNA 1 (targeting the first exon) and gRNA 2 (targeting the fourth exon) are highlighted in purple, PAM sequence is marked with a green rectangle, Cas9 cut site is indicated by a black arrow. Recognition sequences for the restriction enzymes (SalI and HpyF10VI) are marked in light grey; blue arrows indicate position of cut sites.

Both gRNA target sites contain restriction enzyme recognition sites, which facilitate the detection of resulting mutations (Fig. 2). We chose the vector pDIRECT_22C designed for CRISPR-Cas9 editing in plants, which allows efficient assembly of multiple gRNA spacers in a single step. To generate stably transformed plants, we employed *Agrobacterium*-mediated transformation of leaf explants followed by *de novo* organogenesis. We obtained 50 transgenic lines for Désirée. The regeneration in cv. Otava was much slower and less effective, as it showed delayed and less prominent callus formation and reduced shoot regeneration. Thus, we obtained only one transgene-positive line. The obtained transgenic lines are further referred to as the T0 generation.

#### 2.1.3. Evaluation of CRISPR-Cas9 Induced Mutations in T0 Generation

The identification of mutations requires an approach enabling to separately analyse all allelic variants due to tetraploid nature of potato genome and potential mosaicism in regenerated individuals. For initial screening of T0 generation, we used a combination of cleaved amplified polymorphic sequences (CAPS) analysis of the PCR products using primers annealing to all *BEL5* alleles. In our case, the CAPS analysis was applicable for both Cas9 cleavage sites as both contain restriction enzyme recognition sites (Fig. 2). Sequence edits at these sites disrupt restriction enzyme recognition sequences and prevent cleavage. The results of CAPS analysis for cv. Désirée are provided in Fig. S2, and editing efficiency is summarized in Fig. 3. Mostly, we observed a mixture of edited and non-edited sequences for both gRNA target sites indicating that the edits were present only in some alleles or a subset of cells. Interestingly, in individual lines the editing efficiency differed for particular target site (Fig. 3B). We found only one line with a completely edited target site at exon 4 (gRNA2 site; line 60). In addition to short indels that prevailed, we also observed larger insertions/deletions in some lines, including line 60, as evidenced by products of unexpected lengths apparent on the electrophoretic gel (Fig. S2). In Otava, we did not detect any edits at either gRNA target site in the only positively genotyped line (Fig. S3), so we omitted this cultivar from further analyses.

**Fig. 3:**
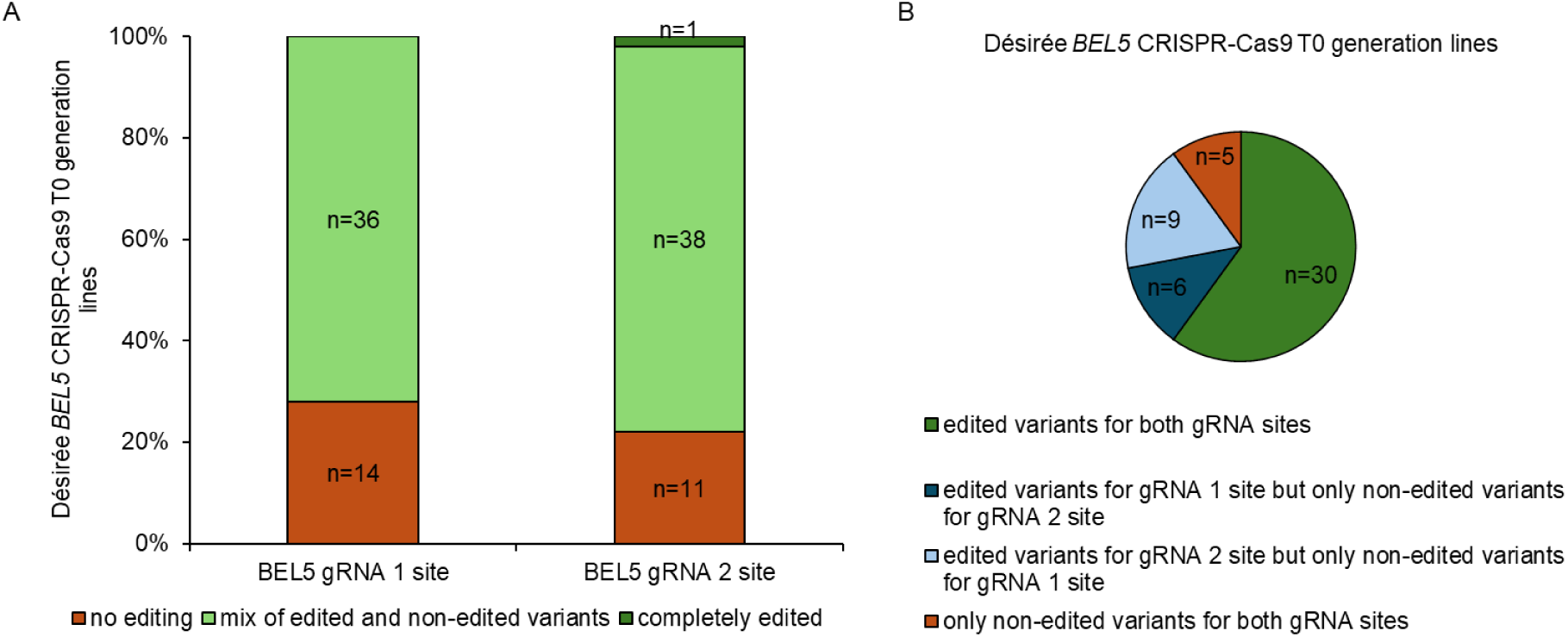
Editing efficiency of *BEL5* in lines of T0 generation of cv. Desirée. (A) Proportion and number of regenerants showing no-editing/ mixture of edited and non-edited variants/complete editing at *BEL5* gRNA 1 site and gRNA 2 site. (B) Representation of lines with: edited variants for both gRNA sites, only one gRNA site, and no gRNA site (here, edited variants represent all lines where at least some of the alleles were mutated). Analysis is based on CAPS screening (Fig. S2).

Based on the CAPS screening, we selected candidate transgenic lines for evaluation of mutations by Sanger sequencing of PCR amplicons enclosing individual gRNA target sites (Fig. S1). The sequence data underwent TIDE software analysis, which allowed to analyse mixed sequences (different alleles/potentially mosaic) and to estimate sequence changes, thereby to predict whether the gene function was disrupted. We excluded lines with short in-frame indels that were unlikely to result in a gene knockout from subsequent analyses. In four lines (line 60, 65, 89, and 91), we detected high proportion of short frameshift and/or long indels. A summary of the detected mutations is presented in Fig. 4A, graphical outputs from the TIDE analysis are provided in Fig. S4. For short indel frameshift mutations, the estimated efficiency ranged from 44 % to 74 % at the gRNA 1 site and from 23 % to 98 % at the gRNA 2 site. Moreover, we confirmed the presence of a more complex mutation resulting in a longer PCR amplicon at the gRNA 1 site in the line 60. The highest editing frequency was estimated for the line 60 at the gRNA 2 site (98%), which was the only one that we considered completely edited, as no non-edited sequence was detected by CASP (Fig. S2B) and TIDE (Fig. S4) analyses. We also examined the occurrence of mosaicism in the line 60, as it may affect overall frequency of edits. We observed highly similar mutation patterns at both gRNA target sites across six consecutive leaves, covering all potential variants within the potato spiral phyllotaxy. The results indicate absent or negligible sectorial mosaicism (Fig. S5).

**Fig. 4:**
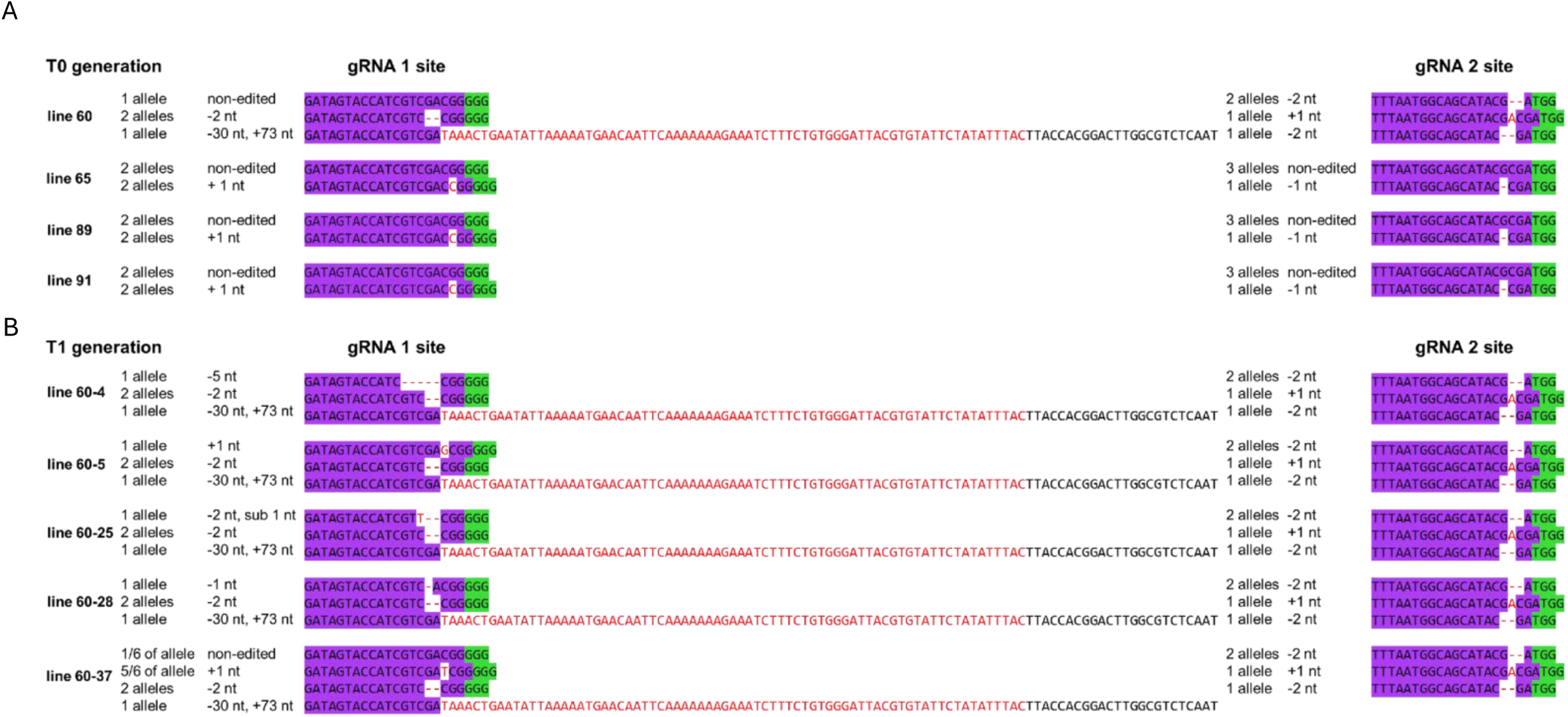
CRISPR-Cas9 induced mutations in selected BEL5 mutant lines of Désirée: (A) T0 generation, (B) T1 generation. gRNA target sites are shown in purple, PAM sequences in green, and CRISPR-Cas9-induced edits in red. Mutations were first identified by CAPS (see Figs. S2 and S7) and then validated by Sanger sequencing combined with TIDE analysis (Figs. S4 and S8). Line 60 (T0 generation) and its T1 progeny (60-4, 60-5, 60-25, 60-28 and 60-37) were also analyzed by Nanopore sequencing of the respective PCR amplicons (the sequence data are available in the Zenodo repository). Allelic composition was determined based on the percentage distribution estimated by TIDE analysis, and results of amplicon sequencing.

#### 2.1.4. Repeated *de novo* regeneration promoted the emergence of new mutations and thereby facilitated the achievement of a complete knockout

The previously unedited sites may still be targeted thus we first verified whether *Cas9* remained expressed in the four selected T0 lines (60, 65, 89, and 91) (Fig. S6). To get rid of potential periclinal mosaicism we performed an additional round of *de novo* regeneration from leaf explants. As dedifferentiation and cell cycle progression are associated with dynamic chromatin structure changes, we also expected potential increase in target-site accessibility facilitating additional editing. Indeed, CAPS analysis of the gRNA 1 site in several tens of regenerated descendants of each line (referred collectively to as the T1 generation) clearly indicated an increased proportion of edited sequences (Fig. S7). However, only among the line 60 descendants, we could identify five lines (designated 60-4, 60-5, 60-25, 60-28, 60-37) that completely lacked the digested product (Fig. S7A), indicating full editing, and similar outcomes were obtained by Sanger sequencing followed by TIDE analysis (Fig. S8). All descendants of the line 60 inherited the large mutation from the progenitor line (Fig. S7A). Additionally, similarly as in the progenitor line 60, we detected the 2-nt deletion in the five T1-descendants at the frequency expected for two alleles. Interestingly, the presumed one allele, which remained unedited in the progenitor line 60, underwent distinct edits in individual descendant lines. We identified these mutations as short frameshift indels (Fig. S8). The gRNA2 site seemed completely edited already in the progenitor line 60, and it was kept in the progeny lines, as we confirmed both based on CAPS results (Fig. S7E) and Sanger sequencing followed by TIDE analysis (Fig. S8).

To verify the identified mutations and precisely assess their allelic distribution, we performed amplicon sequencing of the entire *BEL5* gene and its flanking UTRs, using Oxford Nanopore Technologies in the potentially fully edited lines (raw sequence data available in the Zenodo repository; summarized in Fig. 4B). This analysis confirmed mutations previously detected by CAPS and Sanger sequencing followed by TIDE analysis. Additionally, high read depth of this approach, enabled us to confirm *de novo* origin of additional mutations in the T1 progeny, since they were not detected in the progenitor line 60. We were also able to characterize a previously detected long mutation at the gRNA1 target site, which was found to consist of a 30-nt deletion coupled with a 73-nt insertion. Interestingly, the inserted 73-nt fragment showed an exact match with potato mitochondrial sequence (GenBank: MN104803.1). Moreover, in line 60-37, the analysis revealed a minor proportion of unedited sequence at the gRNA1 target site (approximately one-sixth of allele 1), which escaped detection by previous methods. The presence of this very small portion of unedited sequences may indicate a possible mosaicism in this line, albeit represented in minute proportion.

Based on the assignment of mutations to specific alleles, we were able to assemble the predicted amino acid sequences according to the whole sequence of each *BEL5* allele (Fig. S9). We revealed the presence of premature stop codons already in the first exon for all alleles of lines 60-4, 60-5, 60-25, and 60-28, indicating a complete knockout of *BEL5*. Although most sequences also carried stop codons in the first exon in the line 60-37, a small proportion of them contained a premature stop codon solely in the fourth exon, which might not ensure a complete inactivation of the BEL5 function. Therefore, this line was excluded from subsequent phenotypic analyses.

### 2.2. Phenotypic analysis: *BEL5* knockout does not prevent tuberization but causes slightly delayed tuber onset

In WT and *BEL5* knockout lines 60-4, 60-5, 60-25, and 60-28, tuberization was monitored weekly up to 35 days after transfer (DAT) to *ex vitro* tuber inductive short day (SD) hydroponic conditions. Whereas tuberization was initiated in all WT plants already 28 DAT, the frequency of tuberizing plants was significantly lower in the mutant lines (20-80%; Fig. 5A). Also, the number of tubers per plant was significantly reduced in the mutants at 28 DAT (Fig. 5B). By 35 DAT, these differences diminished, all mutant lines except line 60-25 gradually reached the same number of tubers, which continued to grow until they were comparable in size to WT (Fig. 5C). Importantly, *BEL5* knockout did not significantly change whole-plant biomass (Fig. 5D, E). We further evaluated overall tuber yields under natural-like conditions (soil-grown plants, under long day (LD) photoperiod, harvested at the end of the growing period, 110 DAT) and observed no significant impact on either total yield or tuber number (Fig. 6). Taken together, our data indicate that BEL5 accelerates the onset of tuberization but is not essential for further tuber development.

**Fig. 5:**
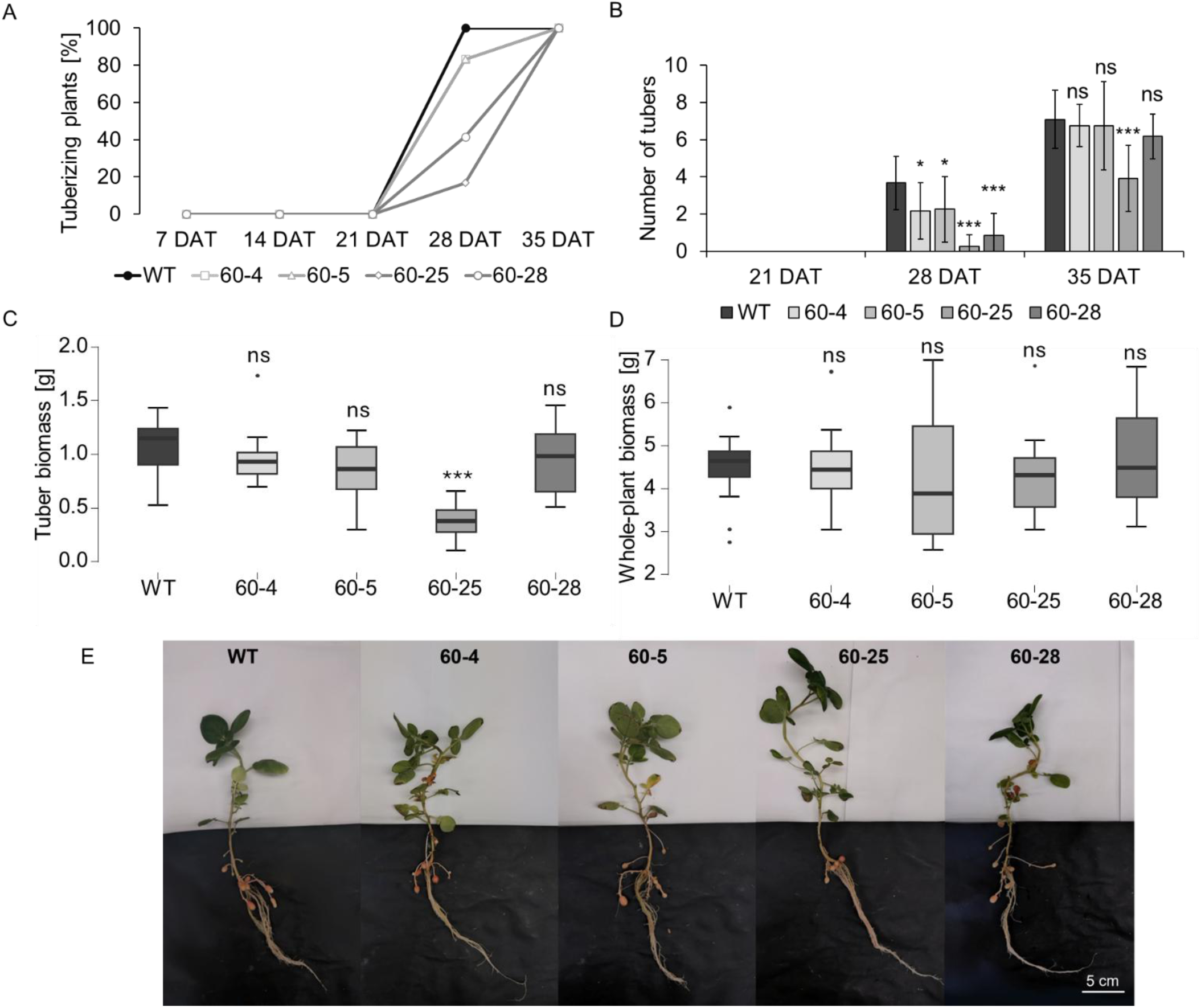
Timing of tuberization onset and growth parameters in selected lines with complete *BEL5* knockout: All parameters were evaluated in SD hydroponically grown plants at 35 DAT except (A) timing of tuber onset and (B) tuber number that were followed weekly. (A) Mutant plants show delayed tuberization. (B) Average number of tubers was significantly lower in mutants at 28 DAT, but this difference started vanishing by 35 DAT. (C) Average tuber biomass was comparable to WT in all transgenic lines except line 60-25. (D) Average whole-plant biomass was not affected in mutants. (E) Appearance of tuberizing plants. WT Désirée and four *BEL5* knockout lines (60-4, 60-5, 60-25, 60-28); n=12; for (B) error bars represent the standard deviations. ANOVA-One-Way Analysis of Variance, Dunnett’s Two-Sided Multiple-Comparison Test With Control was used for statistical evaluation; asterisks indicate statistically significant differences *** (α = 0.001); * (α = 0.05); ns (not significant).

**Fig. 6.**
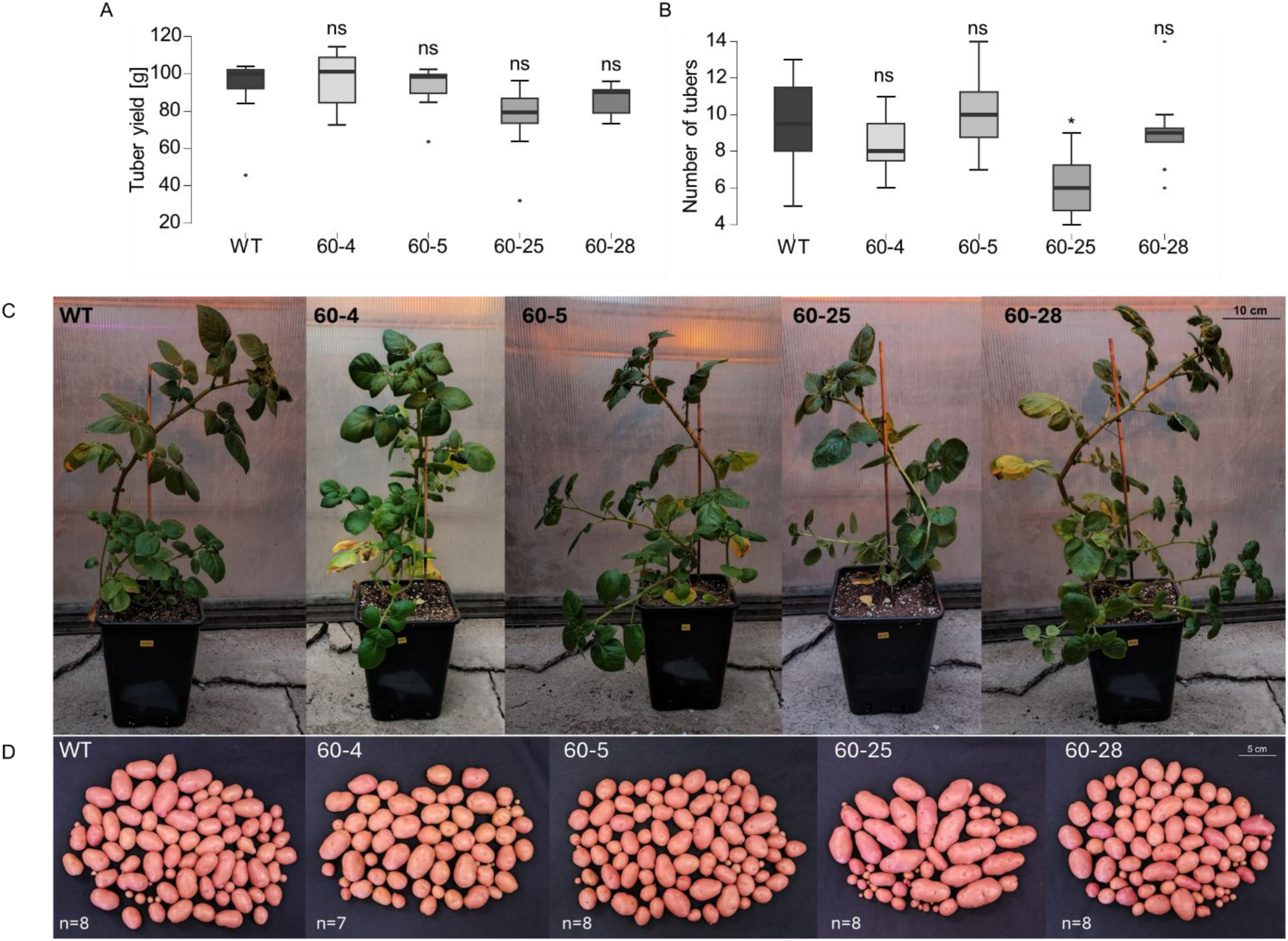
Overall tuber yield in selected lines with complete *BEL5* knockout. (A) Average tuber yield and (B) average number of tubers harvested at the end of the growing period (110 DAT) showed no systematic difference compared to WT. (C) Plant appearance at 60 DAT and (D) appearance of tubers harvested at 110 DAT. Soil-cultivated plants under LD photoperiod in the greenhouse; WT Désirée and four BEL5 knock-out lines (60-4, 60-5, 60-25, 60-28); n=7-8; ANOVA-One-Way Analysis of Variance, Dunnett’s Two-Sided Multiple-Comparison Test With Control was used for statistical evaluation; asterisks indicate statistically significant difference * (α = 0.05); ns (not significant).

## 3. Discussion

The modern cultivated potato *Solanum tuberosum* ssp. *tuberosum* has a highly heterozygous tetraploid genome, resulting in complex segregation of alleles that complicates sexual breeding and efficient selection of desirable traits. Moreover, most cultivars exhibit reduced fertility. As a result, cultivated potatoes are predominantly propagated vegetatively. These biological characteristics make conventional breeding a long-term and challenging process (Bradshaw, 2022). In recent years, the introduction of the CRISPR-Cas9 genome-editing system has revolutionized plant breeding by enabling precise modification of specific genes associated with key agronomic traits, thereby accelerating the breeding process (Tuncel *et al*., 2025). Although this approach offers considerable advantages over conventional breeding, its application in cultivated potato is not as straightforward as in model diploid species with sexual reproduction and, thus, requires careful optimization and balancing of several key factors. In this study, we present a workflow for achieving CRISPR-Cas9 induced complete gene knockout in tetraploid cultivated potato, with broader applicability to other vegetatively propagated polyploid species. Using *BEL5* gene as a target, we optimized key steps and evaluated various approaches for detection of induced mutations to effectively derive a complete gene knockout in modern potato and performed basic phenotypic characterisation of mutants affected in this key transcription factor involved in the tuberization signalling.

### 3.1. Key parameters for successful gene knockout in potato: challenges and optimization

Careful selection of the genotype is crucial for successful gene editing. Due to potential differences in regulatory signalling pathways the insights from model genotypes may not be fully accommodated to evolutionary distant ones (Hardigan *et al*., 2017). Considering also the availability of sequencing data, we chose two potato cultivars Otava and Désirée (Sevestre *et al*., 2020; Sun *et al*., 2022). These varieties differ in timing of tuber onset and thus their comparison may help to specify the BEL5 role in tuber initiation and subsequent tuber development and filling. We also included a list of genotypes with fully sequenced genomes that could be considered suitable for CRISPR-Cas9 applications, including some cultivated varieties (see Table S1).

For initial gRNA design we used CRISPR-P 2.0 that is developed to streamline gRNA design for diverse Cas systems with varying PAM motifs, allowing to predict on-target efficiency and potential off-targets (Liu *et al*., 2017). Alternative softwares are also at disposal (Concordet and Haeussler, 2018; Naito *et al*., 2015). We verified that the selected gRNAs matched all *BEL5* alleles in our cultivars and had minimal off-target risk. For cv. Otava, analyses were straightforward thanks to the availability of haplotype-resolved genomic data and their integration into the CRISPOR platform (Concordet and Haeussler, 2018; Sun *et al*., 2022). Advanced analyses are required for non-reference genotypes, the gRNA design can be achieved using e.g., CRISPR-Local or Geneious Prime tool, allowing the upload and analysis of custom whole-genome sequencing data (Kearse *et al*., 2012; Sun *et al*., 2019). For Desirée, our analyses were performed using unassembled short Illumina reads (Sevestre *et al*., 2020) thus requiring the additional processing (mapping corresponding to *BEL5* allelic variants to the reference sequence in Geneious Prime, and validation using long-read Oxford Nanopore sequencing). Moreover, the difficulty in predicting off-target effects caused by the large volume of short-read data was addressed by implementing advanced bioinformatic analyses, based on Fu *et al*., (2014) and Zhang *et al*., (2016). As late as in 2025, the haplotype-resolved genome of cv. Désirée was published (Godec *et al*., 2025) and incorporated into CRISPOR thus we could employ it only for final confirmation of accuracy in allele assemblies (see Table S2) and potential off-targets.

Beyond selecting gRNAs with high specificity that match all the alleles, it is important to consider additional parameters in order to achieve a complete gene knockout (Přibylová and Fischer, 2024). Our effort was to select a gRNA targeting the coding sequence near the start codon that increases the likelihood of generating early frameshift mutations. Besides a gRNA targeting the first exon of *BEL5* gene we added a gRNA targeting the fourth exon as multiple gRNAs per gene can further enhance chances to achieve complete knockout. When designing gRNAs, it is worth considering that a strategic design can facilitate subsequent mutation screening. Thus, both gRNAs selected for our study contain a restriction site at the expected Cas9 cleavage site, enabling the employment of CAPS analysis.

Using a web-based tool for streamlined vector selection, we chose pDirect 22C, a T-DNA vector for *Agrobacterium*-mediated transformation, enabling to express multiple gRNAs from a single Csy4-processed polycistronic transcript, and employing type II Cas9 from *Streptococcus pyogenes* (SpCas9) (Čermák *et al*., 2017). SpCas9 is the most widely adopted nuclease in plant genome editing, recognizing the abundant NGG PAM motif showing high activity and specificity and is therefore employed in the majority of available plant genome editing vectors (Čermák *et al*., 2017; Tuncel *et al*., 2025). Although alternative natural and engineered Cas variants with distinct PAM specificities have been applied in plants, these systems are still not fully optimized and routinely used in potato (Chincinska *et al*., 2023; Tuncel *et al*., 2025; Zhan *et al*., 2023).

Efficient plant transformation remains among the major limiting factors in the advancement of plant genome editing. In potato, various methods have been employed for both stable and transient transformation, each with its own pros and cons. Transient CRISPR-Cas9 expression (using biolistics or virus-induced gene editing) is sufficient to induce mutations into a target gene (reviewed by Tuncel *et al*., 2025). This approach has the advantage of potentially obtaining “transgene-free” edited lines; however, achieving complete editing throughout the entire plant remains a major challenge as these methods typically affect only a subset of cells. In sexually propagated species, mutations can be induced in germline cells or their precursors, allowing potential production of fully edited progeny (Hamada *et al*., 2018; Lee *et al*., 2025). In vegetatively propagated species such as potato, alternative strategies include tissue culture-based regeneration from edited somatic cells or *de novo* ectopic induction of meristems *in planta* (Maher *et al*., 2020; Tuncel *et al*., 2025). Nevertheless, these methods carry the risk of producing mosaic plants (Bánfalvi *et al*., 2020; Maher *et al*., 2020). To address these challenges, a protocol employing transient CRISPR-Cas9 expression in protoplasts followed by *de novo* organogenesis—yielding fully edited plants without transgene integration—was developed (Johansen *et al*., 2019). However, plants regenerated from protoplasts have been shown to exhibit large-scale genome instability, including aneuploidy and structural chromosomal alterations and therefore these approaches cannot be considered as an optimal solution (Fossi *et al*., 2019). Stable potato transformation – typically mediated via *Agrobacterium tumefaciens* or *Rhizobium rhizogenes*, albeit requiring tissue culture-based *de novo* regeneration (potentially producing mosaic plants), carries a significantly lower risk of undesired genomic alterations (Butler *et al*., 2020; Fossi *et al*., 2019).

Considering the factors mentioned above, we employed *Agrobacterium*-mediated transformation of leaf explants, followed by *de novo* organogenesis, to generate stably transformed plants. Intriguingly, genotype-dependent regeneration potential emerged as a significantly limiting factor. We were able to obtain a sufficient number of transgenic lines for Désirée, but not for Otava. It is in accordance with (Park *et al*., 2023) who showed a high regeneration potential and transformability of cultivar Désirée. However, no reports in this regard exist for cv. Otava, highlighting the need to optimize the transformation protocol for each cultivar. The majority of regenerated transgenic lines of cv. Désirée exhibited some editing, with 60% exhibiting mutations at both gRNA target sites, 30% at a single site, and only 10% lacking CRISPR-Cas9–induced mutations at either target locus. However, most edits— apart from one gRNA site in a single line— were incomplete in the T0 generation (not all alleles or cells carried an edit). In this context, an integrated transgene can be advantageous; if the CRISPR–Cas9 system remains active it can repeatedly target the desired locus, thereby potentially increasing editing efficiency (Přibylová and Fischer, 2024). Notably, continuous CRISPR–Cas9 expression is not inherently associated with elevated off-target effects across generations when highly specific gRNAs are employed (Impens *et al*., 2022).

Since selfing or crossing to inherit edited alleles is not feasible in potato due to already mentioned challenges in sexual reproduction, exploiting continuous CRISPR-Cas9 activity provides an alternative strategy to achieve edits in all alleles. Given that CRISPR–Cas9 efficacy is drastically influenced by epigenetic features of the target locus (Přibylová *et al*., 2022; Weiss *et al*., 2022), we performed a second round of *de novo* regeneration, since this process is associated with dynamic epigenetic reprogramming and extensive chromatin remodelling (Lee and Seo, 2018), and therefore may improve DNA accessibility and editing. Previous studies in other species have shown that repeated *de novo* regeneration can also reduce mosaicism through reselection of transgenic cells (Awasthi *et al*., 2023; Ding *et al*., 2020; Li *et al*., 2023; Malabarba *et al*., 2020). However, we found that it can also promote the emergence of completely new CRI SPR-Cas9 edits in the T1 progeny that were not present in the initial T0 line. This phenomenon has not yet been described. Given that *Cas9* remained expressed and mosaicism was negligible, the newly mutated alleles nearly certainly arose in connection with the second round of *de novo* regeneration. Thus, repeated *de novo* regeneration proved as an effective strategy for increasing the likelihood of achieving full-allelic editing, applicable even in vegetatively propagated polyploid species.

A logical future step would be the generation of fully edited, transgene-free potato lines. Unlike sexually propagated species, in cultivated potato, CRISPR–Cas9 cassette cannot be easily segregated out in subsequent generations (Hamada *et al*., 2018; Lee *et al*., 2025). One potential strategy to induce excision of the integrated transgene cassette, as demonstrated e.g. in apple (Pompili *et al*., 2020). Alternatively, repeated transient expression of *Cas9* and gRNAs could be employed to target unedited alleles in successive rounds, thereby increasing the probability of achieving complete knockout while avoiding stable transgene integration. Nevertheless, repeated virus infections or successive biolistic deliveries can expose plant cells to additional stress and make the overall process of complete knockout preparation more complicated.

Another significant difficulty in polyploid, vegetatively propagated, and potentially mosaic plants, such as cultivated potato, is the precise and reliable detection of allele-specific CRISPR–Cas9-induced mutations and verification that the plant is fully edited. A wide range of methods is available, but each has its own limitations. In practice, a combination of initial rapid assays for screening and high-resolution sequencing-based methods for validation is advisable. To facilitate future work in this area, we have compiled a summary table of detection methods used in polyploid species, together with an overview of their applicability (Table S4).

In our case, we screened our transgenic lines using CAPS analysis. This method is simple, rapid and suitable for screening large set of samples. It is sensitive even to one-nucleotide indels and allows estimation of the portion of edited DNA in the sample. Due to the presence of additional PCR products of unexpected lengths on the electrophoretic gel, we were also able to detect larger insertions/deletions, likely caused by some of error-prone DNA repair pathways, possibly polymerase θ–mediated end joining (Kamoen *et al*., 2025). However, CAPS analysis does not provide information on the exact sequence edits in the target sequence. To initially characterize individual edits, we selected Sanger sequencing of amplicons encompassing individual target sites using primers ensuring detection of all *BEL5* alleles. Subsequent analysis using a specialized TIDE software, is advantageous due to the capability of interpreting overlapping sequence traces (Brinkman *et al*., 2014). This approach provided a valuable preliminary assessment of editing outcomes. Alternatively, sequencing of allele-specific amplicons may be employed but it requires reliable sequence data of all haplotypes. However, for final validation of gene knockout in selected lines, an accurate next-generation amplicon sequencing is highly recommended. We utilized Oxford Nanopore technology, generating thousands of reads per amplicon, to get the complete allele-resolved sequence of *BEL5* gene. This high sequencing depth enabled us not only confirmation of previously identified mutations but also the detection of a low portion of one allele (designated allele 1) that remained unedited in the line 60-37 that had appeared fully edited using earlier methods, but it may retain a minimal mosaicism.

Based on our results, we strongly recommend not relying exclusively on Sanger sequencing—either applied to mixed samples covering all alleles with subsequent software analysis, or to allele-specific PCR products, as well as on a PCR screen of even bigger number of bacterial colonies — as the only method for confirming a complete editing. These approaches, however, still routinely applied in published studies, lack sufficient sensitivity to detect a small portion of unedited alleles in mosaic plants. NGS amplicon sequencing has proven to be the most sensitive and accurate method for this purpose, however, given its higher cost, it is more appropriate to use it only for the final approval stage. In our case, the obtained sufficient sequencing depth using long reads covering the entire *BEL5* gene, enabled us also to accurately resolve edits present in individual alleles. This allowed to translate each edited allele into the amino acid sequence and estimate position(s) of the premature stop codon in each open reading frame. Due to high genome variability and heterozygosity in potato, a careful allele-specific prediction for each genotype should be made for accurate assessment of the functional outcome of gene edits. Surprisingly, this final step is often overlooked in the literature.

### 3.2. Effect of *BEL5* knockout on phenotype

BEL5 has been identified as a key component of potato tuberization regulatory network, positively affecting tuber onset and overall tuber yield (Chen *et al*., 2003; Sharma *et al*., 2016). Nevertheless, functional analyses of BEL5 have been conducted predominantly in transgenic lines with enhanced or reduced expression of the model potato genotype ssp. *andigena* (Banerjee, *et al*., 2006a, b; Banerjee *et al*., 2009; Chen *et al*., 2003; Kondhare *et al*., 2023; Lin *et al*., 2013; Sharma *et al*., 2016). Only a few experiments have addressed BEL5 function in cultivated potato ssp. *tuberosum* cv. Désirée (Cho *et al*., 2015; Kondhare *et al*., 2023). Importantly, in these studies, BEL5 function in cv. Désirée was investigated in relation to Polypyrimidine-tract binding proteins (PTBs), a group of RNA-binding proteins, which, among other functions, promote stability and long-distance transport of *BEL5* transcript. Based on this, the specific role of BEL5 independent of PTBs could not be unambiguously distinguished in these studies.

Here, we report the generation of the first complete and precisely validated *BEL5* knockout in cultivated potato ssp. *tuberosum*, enabling to study the direct effect of BEL5 loss-of-function on tuberization. Interestingly, the mutants exhibited only a slight delay in tuber initiation, yet total tuber yield remained comparable to WT plants. Even though all transgenic lines displayed similar trends, intriguingly, the phenotype of one line was slightly more pronounced. These results reinforce the fact that even in the case of knockout mutants, phenotypic characterization across multiple lines is crucial to rule out potential confounding effects such as epigenetic changes or somaclonal variation (Kaeppler *et al*., 2000).

The less pronounced phenotype, relative to previous reports from ssp*. andigena,* indicates that BEL5 plays a less dominant role in tuberization regulation in this cultivar, or that other factors compensate for its absence. This concept of a genotype-dependent effect of transcription factors related to tuberization is also supported by our recent findings regarding the transcription factor BEL11 (Zounková *et al*., 2026). Based on the initial studies conducted by Ghate *et al*., 2017 in ssp. *andigena*, BEL11 was described as a negative regulator of tuberization. Intriguingly, we showed that in the cultivated potato ssp. *tuberosum* cv. Kamýk, its negative regulatory function appears to be suppressed or even reversed (Zounková *et al*., 2026). Taken together, we suggest that tuberization signals do not act in all-or-nothing manner but rather form a finely tuned network that regulates both the initiation and progression of tuber development. In line with, the absence of BEL5 may be potentially compensated by alternative mechanisms, such as other signalling molecules or modulation of hormonal and carbohydrate pathways (Dutta *et al*., 2024). Given that the BEL5/KNOX and SP6A/FD-like/14-3-3 modules were proposed as two cooperating, partially redundant complexes supporting tuberization, it can be assumed that in the absence of BEL5, SP6A may regulate downstream transcriptional targets to ensure a proper tuber development. (Hannapel *et al*., 2017; Sharma *et al*., 2016). The mutual influence of the intensity of these two signals was observed, for example, in a spontaneously tuberizing potato line (ssp. *tuberosum*, cv. Lada), which exhibited reduced *BEL5* transcript levels but a markedly increased abundance of *SP6A* transcripts (Fischer *et al*., 2008; Ševčíková *et al*., 2017).

Overall, our results highlight the robustness of the tuberization regulatory network in cultivated potato, demonstrating that even when a key component is compromised, the plant may retain the capacity to complete tuber formation with only little alteration from the WT.

## 4. Materials and methods

### 4.1. Plant material

Tetraploid potato (*S. tuberosum* ssp. *tuberosum*), cultivars Désirée and Otava (Table S1) obtained from Potato Research Institute Havlíčkův Brod, Ltd. Czech Republic, were used for the experiments.

### 4.2. Identification of Target Gene Alleles in Selected Potato Cultivars

In the Otava haplotype-resolved genome assembly (Sun *et al*., 2022), four alleles of the *BEL5* gene (Gene IDs: He1-St06G504960; He2-St06G504760; St1-St06G513560 and St2-St06G489680; Table S2) were identified through BLAST analysis, using the annotated *BEL5* sequence (Gene ID: 102577460) from the DM reference genome (The Potato Genome Sequencing Consortium, 2011). For the cultivar Désirée, the sequences corresponding to the four alleles were assembled by mapping Illumina reads (Sevestre *et al*., 2020) available from the NCBI Sequence Read Archive (Acc. No.: PRJNA507597) onto the DM reference genome *BEL5* sequence. Accuracy of assembly was then validated through Nanopore sequencing using Plasmidsaurus service (Premium PCR Sequencing following the provider’s instructions). PCR reactions were prepared using Q5 High-Fidelity DNA Polymerase (New England Biolabs, USA) according to the manufacturer’s instructions. Primers for amplification of complete *BEL5* sequence with an overlap into UTRs (Table S5) were designed to cover all alleles based on Illumina reads assembly. Reads obtained from Oxford nanopore sequencing were then assigned to individual alleles using Geneious Prime software (Kearse *et al*., 2012).

### 4.3. gRNA design and construct assembly

*BEL5* gene specific gRNAs were designed using CRISPR-P 2.0 (Liu *et al*., 2017), available online: http://crispr.hzau.edu.cn/CRISPR2/. Two gRNAs (designated as gRNA 1, targeting the first exon, and gRNA 2 targeting the fourth exon) were selected based on their ability to target all four alleles of both selected potato cultivars (Fig. S1). Potential off-targets were specifically evaluated for each cultivar (Table S3). For Otava, we employed the online tool CRISPOR (Concordet and Haeussler, 2018), https://crispor.gi.ucsc.edu/. For cv. Désirée, dataset (SRR8261493) was screened, with settings allowing up to five mismatches per sequence, employing approaches described in Fu *et al*., (2014) and Zhang *et al*., (2016).

Cloning into plasmid pDIRECT_22C was performed using Golden Gate assembly of PCR products carrying the processing elements, gRNA repeats and target-specific gRNA spacing sequences for Golden Gate junctions, as described in (Čermák *et al*., 2017). The procedure was designed using Webtools for the Voytas Lab Plant Genome Engineering Toolkit (http://crisprmultiplex.cbs.umn.edu/allmaps.php). Primers used for recombinant plasmid assembly are listed in Table S5. The Golden Gate reaction product was transformed into chemically competent *Escherichia coli* DH5α cells using the heat-shock method, followed by cultivation in liquid LB medium (Table S6) for 2 hours at 37 °C. Colonies were selected on solid LB medium (Table S6) supplemented with kanamycin (50 mg/L). Plasmid DNA isolation was done using a miniprep kit (NucleoSpin Plasmid, Mini kit for plasmid DNA, Machery-Nagel) from single colony-derived bacterial culture. Correct assembly was confirmed by Sanger sequencing (Eurofins Genomics; sample preparation followed the provider’s instructions).

### 4.4. *Agrobacterium*-mediated transformation of potato and *de novo* regeneration

Stable transformation of potato plants was achieved using a modified protocol from (Dietze *et al*., 1995). The engineered plasmid was electroporated into *Agrobacterium tumefaciens* strain C58C1, followed by cultivation in liquid YEB recovery medium (Table S6) for 3 h at 28 °C. Colonies were selected on solid LB medium (Table S6) supplemented with kanamycin (50 mg/L) and rifampicin (100 mg/L) (cultivated 3 days at 28 °C). The presence of the transgene was confirmed by PCR using DreamTaq PCR Master Mix (Thermo-Scientific **®**) employing transgene-specific primers (Table S5). A single positive bacterial colony was inoculated into liquid YEB medium (Table S6) supplemented with kanamycin (50 mg/L) and rifampicin (100 mg/L) and cultured overnight at 28 °C to an OD₆₀₀ = 0.6. The resulting bacterial suspension was used for plant transformation. Detached leaves of *in vitro* plants were cut through the midrib and co-cultivated with 50 µL of transformed *A. tumefaciens* suspension in 10 mL of liquid MS medium (Murashige and Skoog, 1962) supplemented with 2% sucrose (w/v) (Table S6), with only the adaxial side of the leaves in contact with the medium, for two days in darkness at 25 °C. The leaves were then dried using sterile filtration paper, transferred to callus induction medium (Table S6), and cultivated for one week at 21°C under LD conditions (16 hours light, 8 hours dark). Afterward, the explants were transferred to the shoot induction medium (Table S6) and cultured with a 7 day – subculture interval under the same conditions for 6 weeks and 13 weeks for Désirée and Otava, respectively. Regenerated shoots were then transferred to solid MS medium supplemented with 3 % sucrose (w/v) (Table S6) and selection antibiotics kanamycin (50 mg/L) and cefotaxime (300 mg/L). The T0 regenerants were further maintained and propagated on MS medium, with subculturing at monthly intervals. For repeated *de novo* regeneration, leaf explants with cuts through the midrib were cultivated one week on callus-inducing medium followed by 6 weeks (with 7 day – subculture interval) on shoot-inducing medium as described above.

### 4.5. Molecular characterisation of CRISPR-Cas9 induced mutations

For verification of the transgene presence in regenerants, genomic DNA was isolated from leaves of *in vitro* grown plants using an optimized rapid DNA isolation protocol (Edwards *et al*., 1991). For the first molecular screening of *BEL5* mutants, evaluation of mosaicism and characterisation of edits after repeated regeneration, DNA was isolated from apical leaf rosette of *in vitro* plants with CTAB method (based on protocol of (Allen *et al*., 2006). PCR reactions were prepared using DreamTaq PCR Master Mix (Thermo-Scientific**®**) following the manufacturer’s protocol either with transgene-specific primers or the primers complementary to all four alleles (Fig. S1, Table S5). For cleaved amplified polymorphic sequences (CAPS) analysis, PCR products of gRNA site 1 and 2 were digested using restriction enzymes SalI and HpyF10VI, respectively (Thermo-Scientific**®**) and analysed in 2,5% TAE agarose gels (Fig. S2, S3 and S7 ). Based on this screen, transgenic lines with high proportion of edited DNA were selected for Sanger sequencing. PCR products for sequencing were prepared in the same way as for CAPS analysis keeping necessary requirements for subsequent software analysis: amplification of all alleles and generation of products of equal length for all alleles. The resulting chromatograms were analysed by Tracking of Indels by DEcomposition (TIDE) software (https://tide.nki.nl/) to reveal sequence type of edits and estimate their proportion (Brinkman *et al*., 2014). Selected lines that appeared to be fully edited at both gRNA target sites were further validated by amplicon sequencing of the entire *BEL5* gene with overlaps to UTRs, using Nanopore sequencing (described in 4.2).

### 4.6. Evaluation of mosaicism

In this study, we adhere to the term ‘mosaicism’ as defined by (Impens *et al*., 2022). Although the term is sometimes used interchangeably with ‘chimerism’, we consider ‘mosaicism’ to be more precise in this context, as the observed genetic differences result from mutations in distinct somatic cells rather than from the presence of cells of different zygotic origins. To evaluate mosaicism in regenerated plants, six consecutive leaves per plant were used, to cover all potential positions in spiral phyllotaxis in potato, as there are typically 3 leaves per 1.5 turns of the spiral, following a 3/5 Fibonacci ratio.

### 4.7. Evaluation of *Cas9* expression

RNA was isolated from leaves of *in vitro* plants using the TRI reagent isolation protocol (Sigma Aldrich**®**). RNA integrity was checked by RNA denaturing agarose gel electrophoresis (Masek *et al*., 2005). To remove contaminating genomic DNA, samples were treated with RNase-Free Turbo DNase (Invitrogen™) according to the manufacturer’s protocol. 1 µg of total RNA from each sample was reverse-transcribed using RevertAid^TM^ reverse transcriptase (Thermo-Scientific**®**) with oligo-dT and RiboLock RNase Inhibitor (Thermo Scientific) following manufacturer’s instructions. Semi-quantitative PCR was performed to estimate the level of the *Cas9* transcript using the primers listed in Table S5. Transcript for *polyubiquitin* (*UBI*) was used as positive reference. Samples without reverse transcriptase and WT were used as negative controls.

### 4.8. Phenotype analysis

#### 4.8.1. Tuberization onset

Parameters related to growth and tuber onset were evaluated in hydroponically cultivated *ex vitro* plants. For this purpose, 4-week-old *in vitro* pre-cultivated plants were transferred to a hydroponic system (Araponics, Liège, Belgium), modified for potato growth. The holders in the lid were modified to allow “underground” shoot (stolon) growth after the *ex vitro* transfer. Roots and a portion of the *in vitro*-arisen shoot (with a consistent number of nodes) were placed in darkness beneath the lid. Each hydroponic system with 1.5 litres of modified ¼ Hoagland solution (Hoagland and Arnon, 1938) contained 12 plants. Plants were cultivated under tuber-inductive conditions: SD (8h light/16h dark), PAR 120 μmol m−2 s−1, white LED supplemented with far-red LED light, and a temperature of 22±2°C for 5 weeks (35 DAT). The percentage of tuberizing plants and the number of tubers per plant were recorded weekly (7, 14, 21, 28, and 35 DAT), all other parameters were determined at 35 DAT.

#### 4.8.2. Overall tuber yields

Overall tuber yields were determined in *ex vitro* soil-grown plants (pre-cultivated 4 weeks *in vitro*) in the greenhouse under LD-adjusted photoperiod, 18±2°C. Plants were cultivated in 7 L pots containing common garden substrate, compost soil, sand and perlite (6:2:1:2). Tubers were harvested at the end of the growing period (110 DAT).

### 4.9. Statistical Analysis

Boxplot graphs were generated using JASP software (Version 0.18.3), line and bar charts were prepared in Microsoft Excel. Error bars represent standard deviations. Statistical analyses were conducted using NCSS 9 software (NCSS, LLC, Kaysville, Utah, USA). A one-way analysis of variance (ANOVA) was performed to evaluate statistical differences. Comparisons between transgenic lines and the WT control were made using Dunnett’s two-sided test (versus control). Statistical significance was determined at P ≤ 0.001, P ≤ 0.01, P ≤ 0.05, and P ≤ 0.1 levels labelled as ***, **, *, and (*), respectively.

## Accession numbers

Illumina reads for cv. Désirée - Sequence Read Archive of the NCBI (PRJNA507597)

NGS-resequencing of cv. Désirée - Sequence Read Archive of the NCBI (SRR8261493)

*BEL5* alleles of cv. Désirée assembled based on Nanopore sequencing, this study – GenBank ID NCBI (PZ132913-PZ132916 for allele 1-4, respectively)

## Author Contributions

AZ and PM conceived the study. AZ, PM and DC performed the experiments. AP conducted the bioinformatic analyses. AZ, PM, DC, AP, DM, JM and TM optimized the experimental methodologies. AZ, PM, DC, AP, LF, CB and FF interpreted the data. AZ wrote the original draft. AZ, PM, DC, AP, TM, LF, CB and FF revised and edited the manuscript. All authors read and approved the final manuscript.

## Acknowledgements

We thank Eliška Kobercová, Lorenzo Mineri, Francesca Giaume, Giulia Ave Bono, and Laura Luoni for their valuable assistance in the laboratory. We also thank Martin Pospíšek for providing access to equipment and facilities. We acknowledge Helena Lipavská for her support.

This work has been funded by a grant TowArds Next GENeration Crops from the ERDF Programme Johannes Amos Comenius under the Ministry of Education, Youth and Sports of the Czech Republic [CZ.02.01.01/00/22_008/0004581].

## Ethics statement

The authors have nothing to report.

## Data availability

The raw datasets including sequencing data are freely available in the Zenodo repository (https://doi.org/10.5281/zenodo.18924477).

## Supporting information Supporting figures

Figure S1: Multiple sequence alignment of cv. Désirée and cv. Otava *BEL5* alleles

Figure S2: CAPS analysis in T0 generation of cv. Désirée

Figure S3: CAPS analysis in T0 generation of cv. Otava

Figure S4: Short indels estimated in T0 generation of cv. Désirée

Figure S5: Evaluation of mosaicism in progenitor line 60 (T0 generation) cv. Désirée

Figure S6: Expression of *Cas9* in selected T0 lines of cv. Désirée Figure S7: CAPS analysis in T1 generation of cv. Désirée

Figure S8: Short indels estimated in T1 generation of cv. Désirée

Figure S9: Amino acid sequences predicted based on individual *BEL5* alleles

## Supporting Tables

Table S1: List of potato varieties with fully sequenced genomes Table S2: *BEL5* alleles in cv. Otava and cv. Désirée

Table S3: Off-target predictions for *BEL5-*targeting gRNAs

Table S4: Commonly used methods for detecting CRISPR-Cas9 mutations in polyploid/mosaic organisms

Table S5: List of primers used in the study

