## Supplementary figures and Tables for "CRISPR-Cas9 Induced Knockout of *BEL5* in Tetraploid Potato: Optimized Methodology via Repeated *de novo* Regeneration and Impact on Tuberization"

**Fig. S1A: Multiple sequence alignment of adjacent sequence region encompassing the gRNA1 target site in *BEL5* alleles of cv. Désirée (see the legend below).**

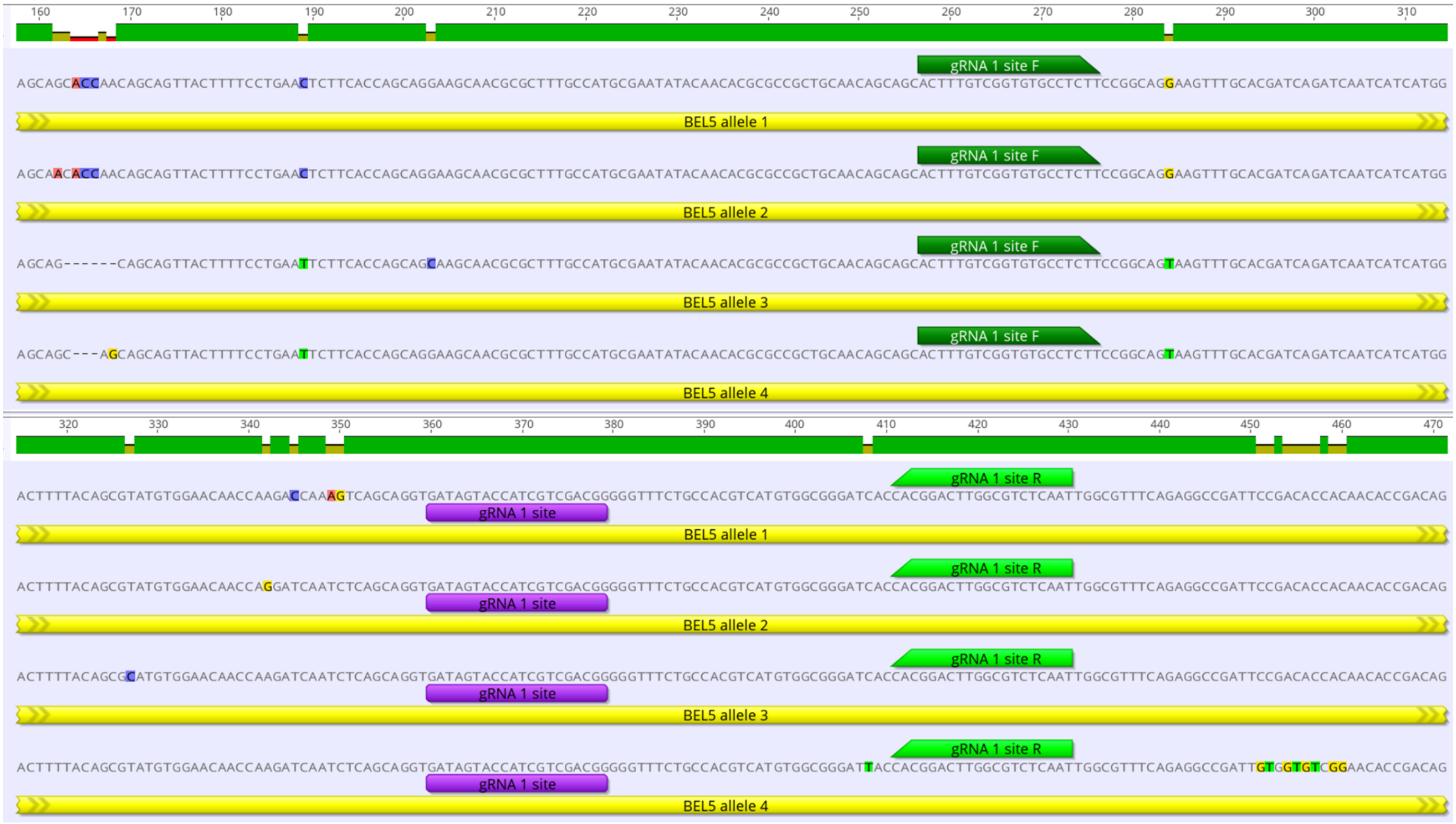

**Fig. S1B: Multiple sequence alignment of adjacent sequence region encompassing the gRNA2 target site in *BEL5* alleles of cv. Désirée (see the legend below).**

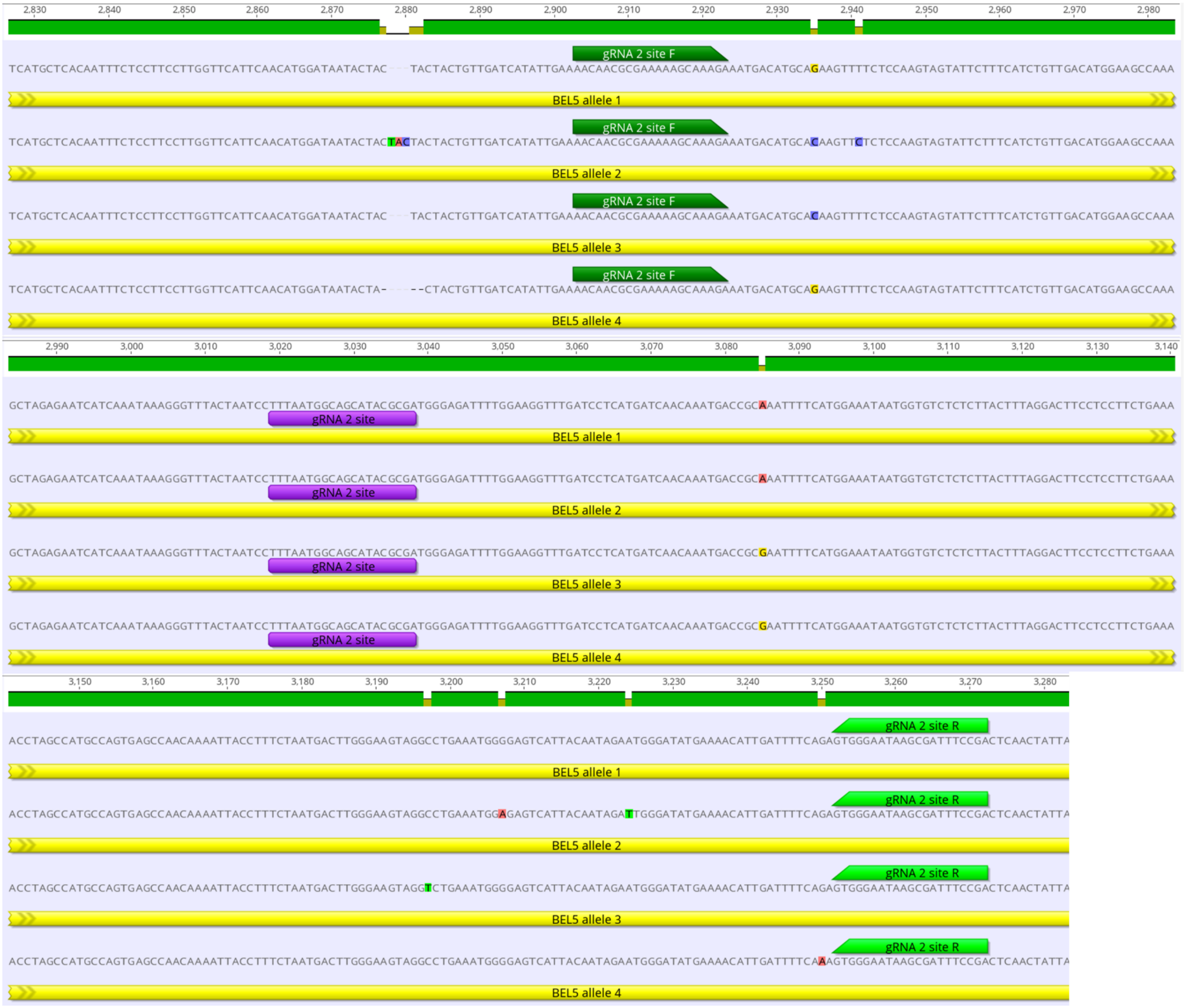

**Fig. S1C: Multiple sequence alignment of adjacent sequence region encompassing the gRNA1 target site in *BEL5* alleles of cv. Otava (see the legend below).**

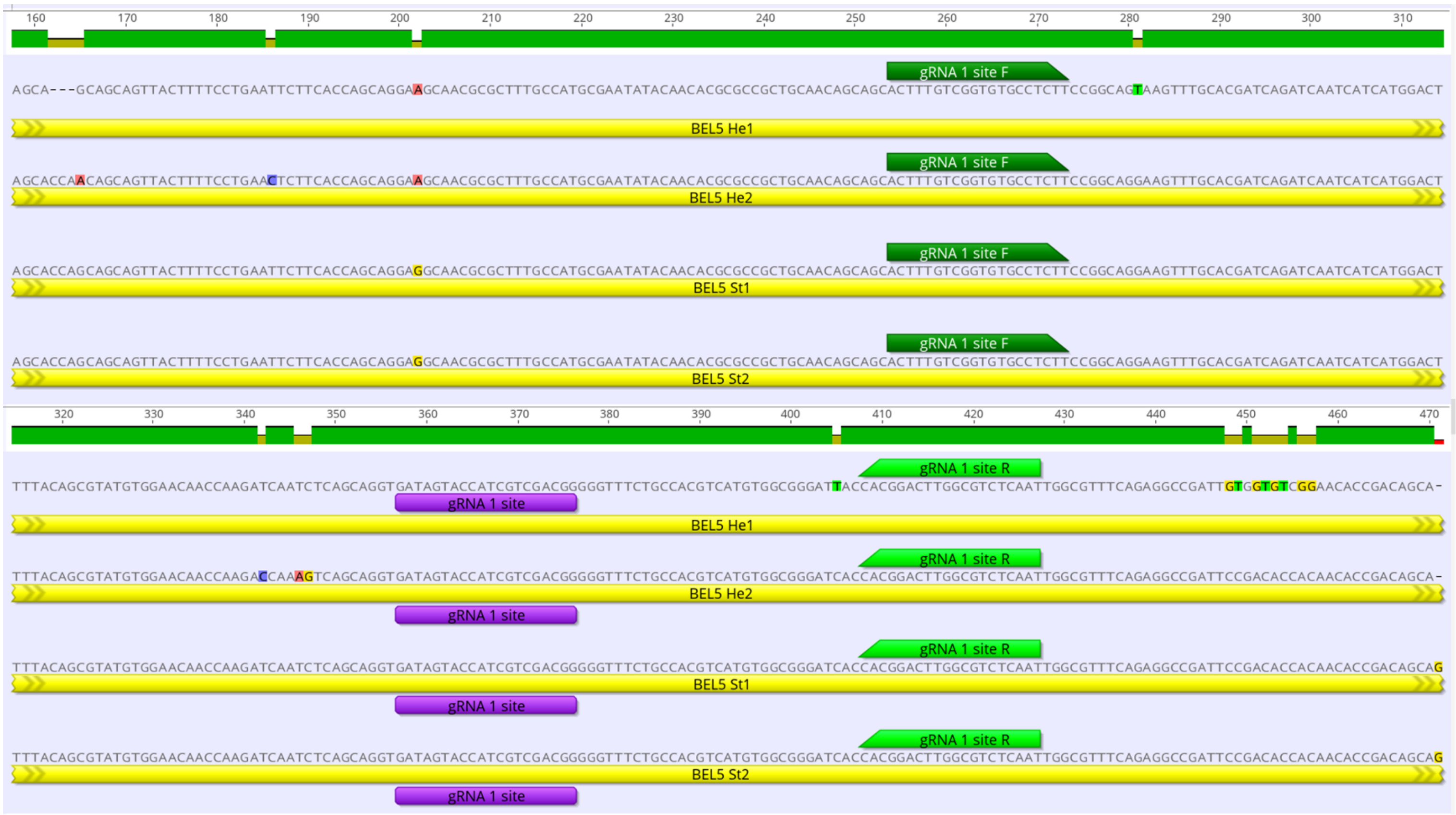

**Fig. S1D: Multiple sequence alignment of adjacent sequence region encompassing the gRNA2 target site in *BEL5* alleles of cv. Otava (see the legend below).**

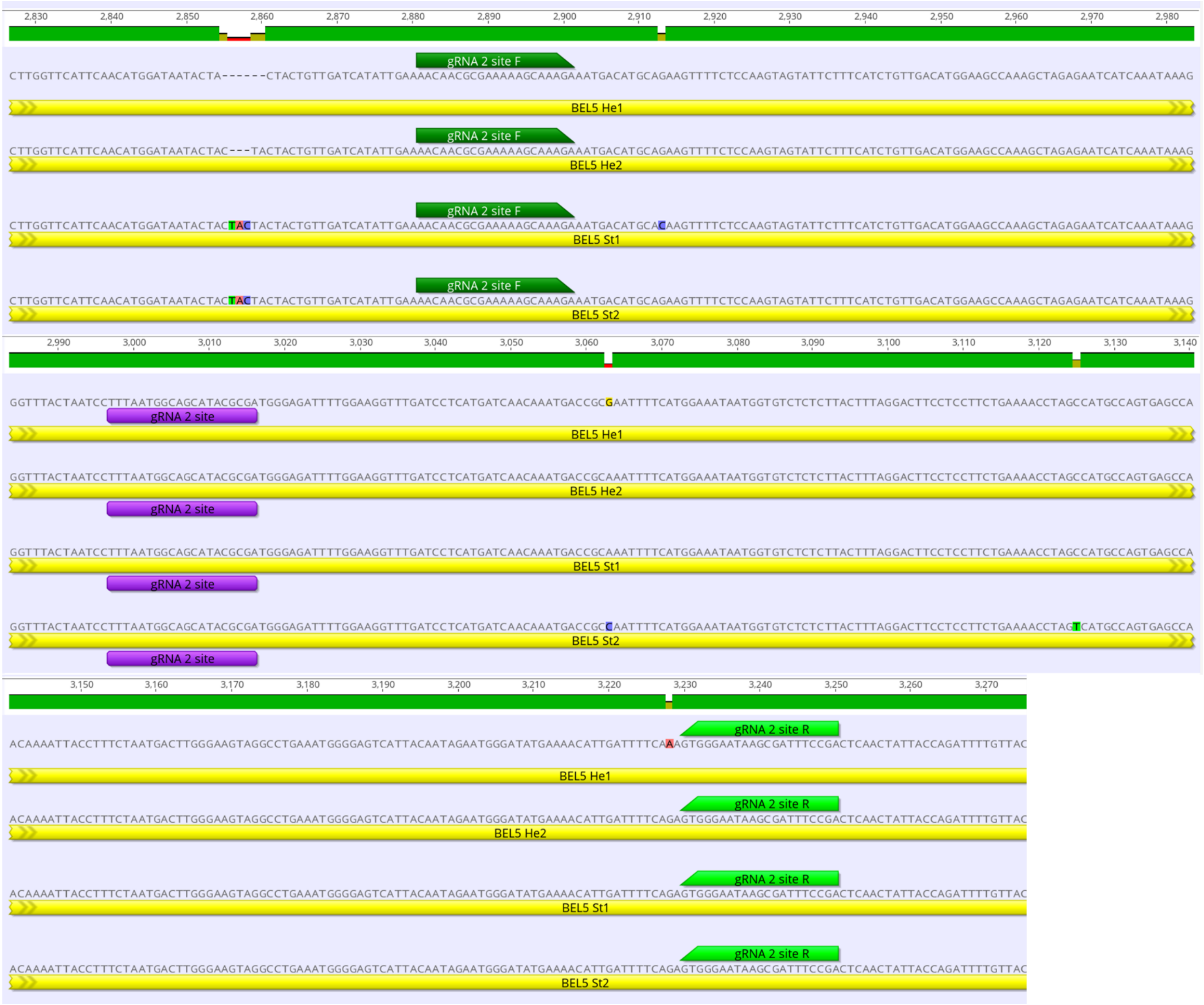


**Fig. S1: Multiple sequence alignment of Désirée (A, B) and Otava (C, D) *BEL5* alleles:** Gene sequences for Otava were obtained from https://spuddb.uga.edu/index.shtml (gene IDs are listed in Table S2). For Désirée, gene sequences were assembled using Illumina sequencing data (downloaded from the NCBI Sequence Read Archive, accession number PRJNA507597) and re-sequenced by Nanopore sequencing (GenBank accession numbers PZ132913-PZ132916 for allele 1-4, respectively). Alignment of the complete *BEL5* coding sequence was performed using the MUSCLE algorithm in Geneious Prime software. Here, we present a partial sequence of the 1st and 4th exon, with highlighted gRNA sites (in purple) and position of primers used for amplification (forward and reverse primers in dark and light green, respectively). PCR products were used for CAPS analysis, Sanger sequencing, and TIDE analysis to detect CRISPR-Cas9-induced mutations.

**Fig. S2A: CAPS analysis in cv. Désirée T0 generation (gRNA 1 target site), see the legend below.**

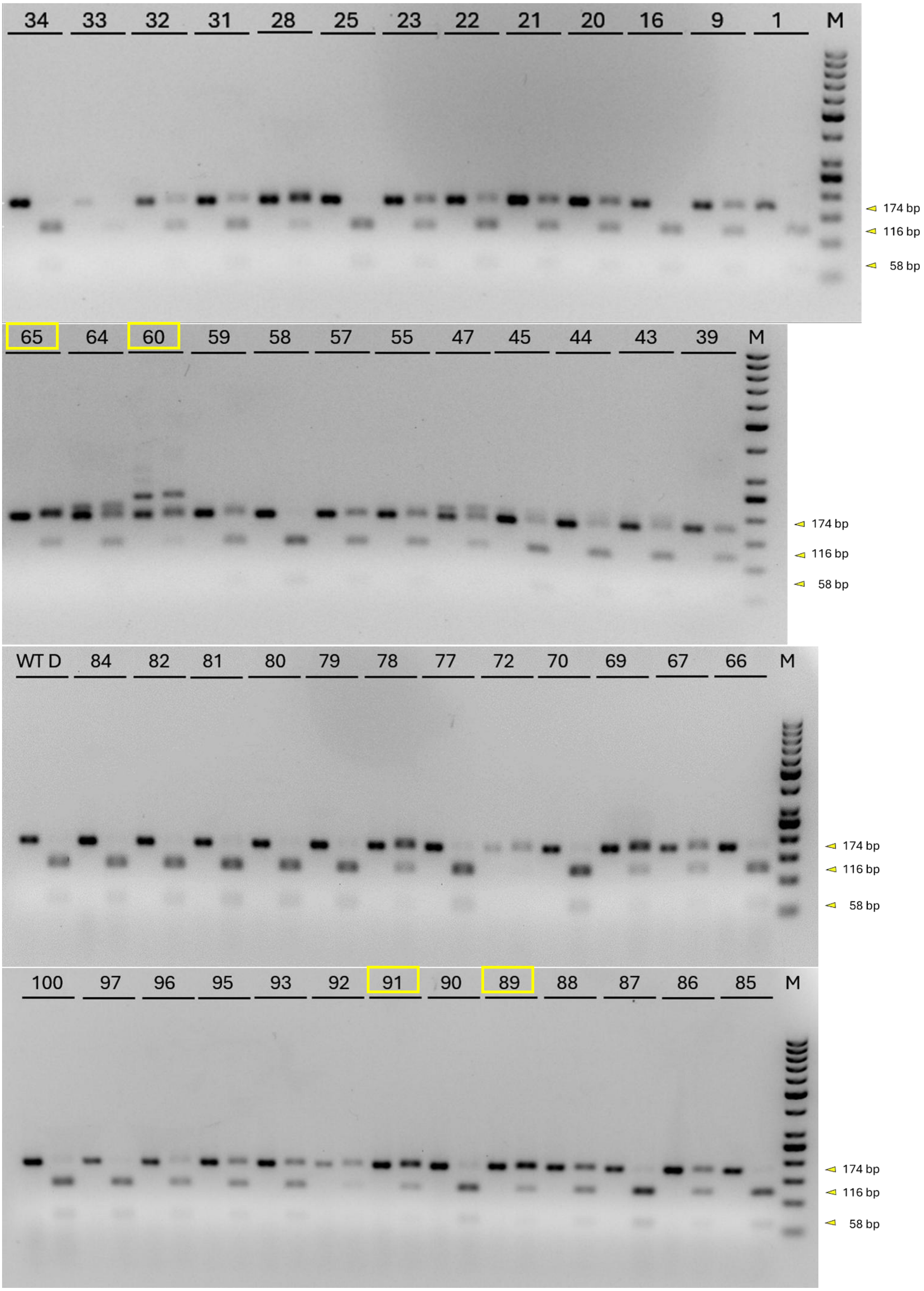

**Fig. S2B: CAPS analysis in cv. Désirée T0 generation (gRNA 2 target site), see the legend below.**

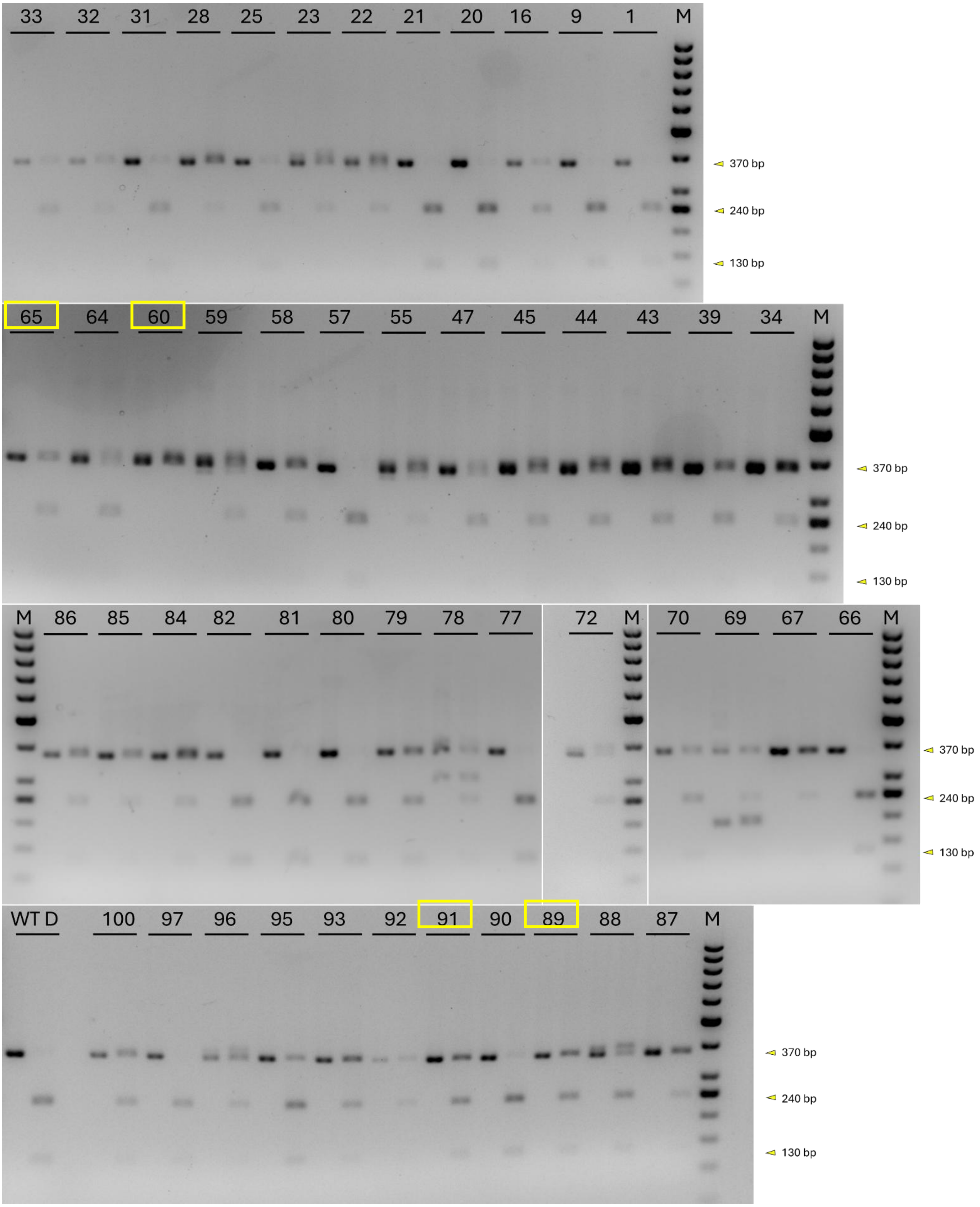

**Fig. S2: CAPS analysis in T0 generation of Désirée:** 50 independent transgenic lines transformed by construct for CRISPR-Cas9 induced knockout of *BEL5* gene were screened for indels using restriction enzyme digestion. M represents size marker (O'GeneRuler 50 bp DNA Ladder (Thermo Scientific). For each transgenic line a pair of samples was tested, from left the first band represents a non-digested control and the second band is a digested sample. WT Désirée (WT D) sample used as positive control showed complete digestion for the SalI site (A) as well as for the HpyF10VI site (B). For (A), transgenic lines 1, 16, 25, 33, 34, 58, 79-82, 84, 85, 97 and 100 showed only digested bands suggesting no editing; lines 9, 20-23, 28, 31, 32, 39, 43-45, 47, 55, 57, 59, 60, 64-67, 69, 70, 72, 77, 78, 86-93, 95 and 96 showed both digested and non-digested bands suggesting a mixture of WT and edited alleles. Moreover lines 47, 60, 64 showed longer product in both digested and non-digested samples suggesting larger mutation. For (B), transgenic lines 1, 9, 20, 21, 57, 66, 77, 80-82, 97 showed only digested bands suggesting no editing; lines 16, 22, 23, 25, 28, 31-34, 39, 43-45, 47, 55, 58, 59, 64, 65, 67, 69, 70, 72, 78, 79, 84-93, 95, 96 and 100 showed both digested and non-digested bands suggesting a mixture of WT and edited alleles. Moreover, in some of these lines we detected also additional bands suggesting longer insertions (lines 23,88 and 96) or deletions (55, 59 and 78); line 60 showed only non-digested band suggesting potential complete editing. The lines marked with a yellow rectangle (60, 65, 89, 91) were selected for repeated *de novo* regeneration.

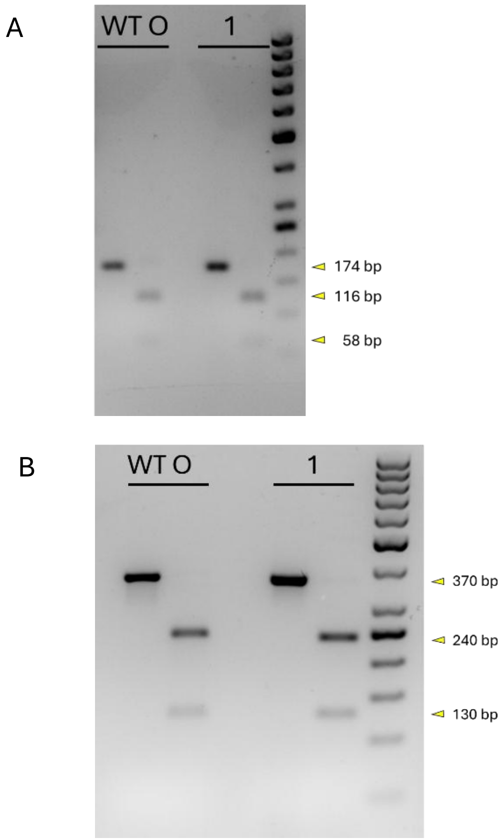


**Fig. S3: CAPS analysis in T0 generation of Otava:** transgenic line transformed by construct for CRISPR-Cas9 induced knockout of *BEL5* gene was screened for indels using restriction enzyme digestion. M represents size marker (O'GeneRuler 50 bp DNA Ladder,Thermo Scientific). For each transgenic line a pair of samples was tested, from left the first band represents a non-digested control and the second band is a digested sample. WT Otava (WT O) sample used as a control showed complete digestion for SalI site (A) as well as HpyF10VI site (B). We detected only digested bands for both gRNA sites in transgenic line 1 suggesting no editing.

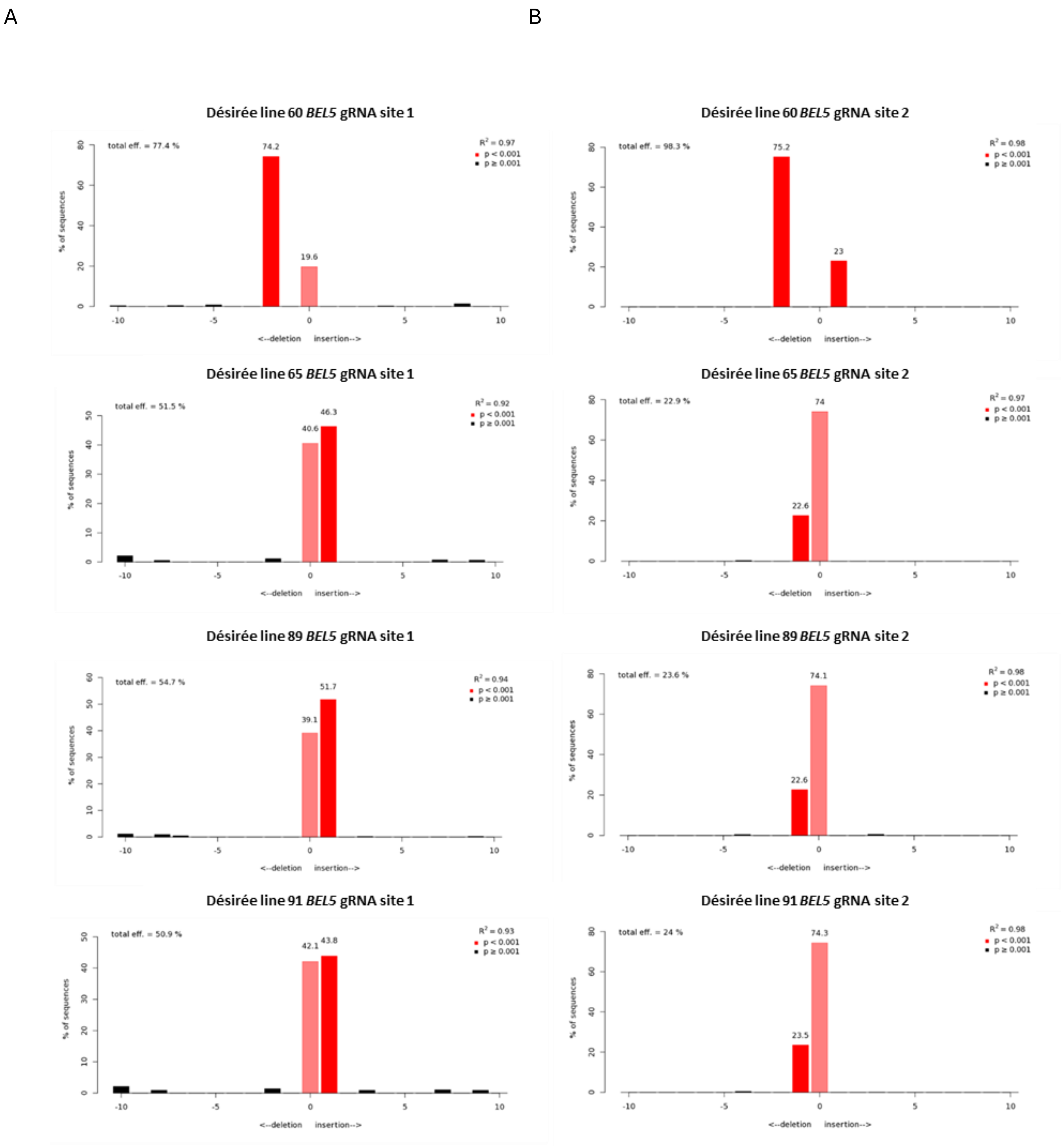


**Fig. S4: Short indels estimated in selected T0 generation lines of cv. Désirée transformed by construct for CRISPR-Cas9 induced knockout of *BEL5* gene:** gRNA 1 site (A), gRNA 2 site (B); lines with higher portion of edited sequences were selected based on CAPS screening (Fig. S2). PCR products covering all alleles were Sanger sequenced and sequence edits and their representation were estimated by TIDE software at gRNA 1 site (A) and gRNA 2 site (B). The obtained results align with the CAPS analysis.

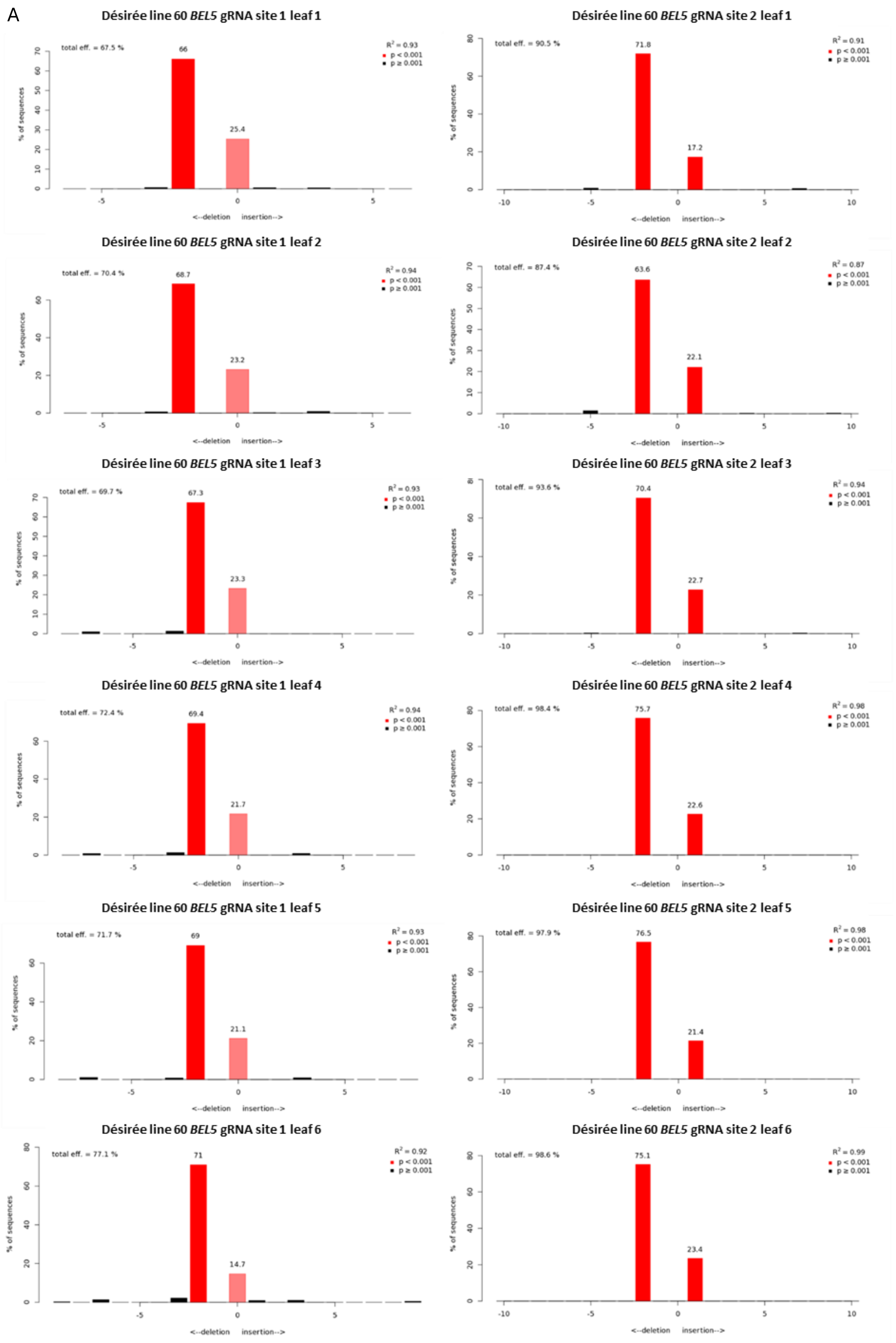
A
Désirée line 60 BEL5 gRNA site 1 leaf 1
total eff. = 67.5 %
66
25.4
R² = 0.93
p < 0.001
p ≥ 0.001
% of sequences
<--deletion insertion-->
Désirée line 60 BEL5 gRNA site 2 leaf 1
total eff. = 90.5 %
71.8
17.2
R² = 0.91
Désirée line 60 BEL5 gRNA site 1 leaf 2
total eff. = 70.4 %
68.7
23.2
R² = 0.94
Désirée line 60 BEL5 gRNA site 2 leaf 2
total eff. = 87.4 %
63.6
22.1
R² = 0.87
Désirée line 60 BEL5 gRNA site 1 leaf 3
total eff. = 69.7 %
67.3
23.3
R² = 0.93
Désirée line 60 BEL5 gRNA site 2 leaf 3
total eff. = 93.6 %
70.4
22.7
R² = 0.94
Désirée line 60 BEL5 gRNA site 1 leaf 4
total eff. = 72.4 %
69.4
21.7
R² = 0.94
Désirée line 60 BEL5 gRNA site 2 leaf 4
total eff. = 98.4 %
75.7
22.6
R² = 0.98
Désirée line 60 BEL5 gRNA site 1 leaf 5
total eff. = 71.7 %
69
21.1
R² = 0.93
Désirée line 60 BEL5 gRNA site 2 leaf 5
total eff. = 97.9 %
76.5
21.4
R² = 0.98
Désirée line 60 BEL5 gRNA site 1 leaf 6
total eff. = 77.1 %
71
14.7
R² = 0.92
Désirée line 60 BEL5 gRNA site 2 leaf 6
total eff. = 98.6 %
75.1
23.4
R² = 0.99

**Fig. S5: Evaluation of mosaicism in progenitor line 60 (T0 generation) cv. Désirée.** PCR products covering all *BEL5* alleles were prepared with DNA isolated separately from six consecutive leaves, Sanger sequenced and short indels and their representation were estimated by TIDE software at gRNA 1 site (A) and gRNA 2 site (B). Similar patterns of mutation representation in individual leaves indicate no or negligible (undetectable by this method) level of mosaicism.

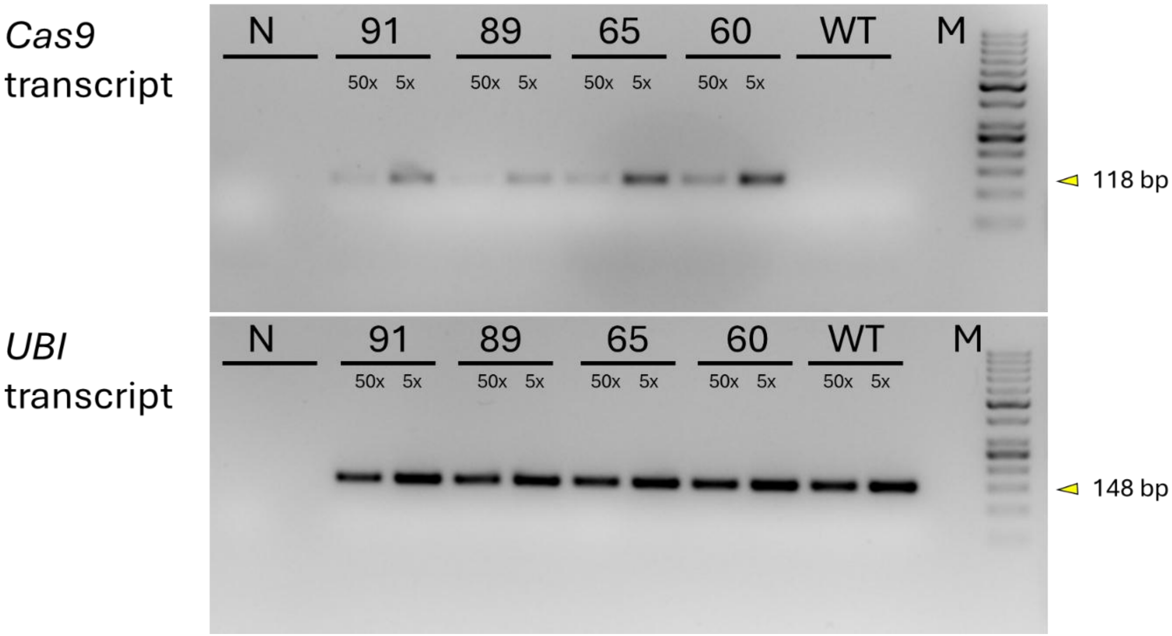


**Fig. S6. Expression of *Cas9* in selected T0 lines:** cv. Désirée transformed by construct for CRISPR-Cas9 induced knockout of *BEL5* gene; *Cas9* transcript was detected in all selected T0 lines (60, 65, 89, and 91), but not in the negative control (N; without reverse transcriptase) or in WT. Amplification of the *polyubiquitin* (*UBI*) transcript served as a positive reference. M represents size marker (O'GeneRuler 50 bp DNA Ladder (Thermo Scientific). RNA was isolated from leaves of 4-week-old, *in vitro*–cultivated plants grown under LD photoperiod. Semi-quantitative RT-PCR was performed using cDNA diluted 5× and 50×.

**Fig. S7A: CAPS analysis in cv. Désirée T1 generation - descendants of line 60 (gRNA 1 target site), see the legend below.**

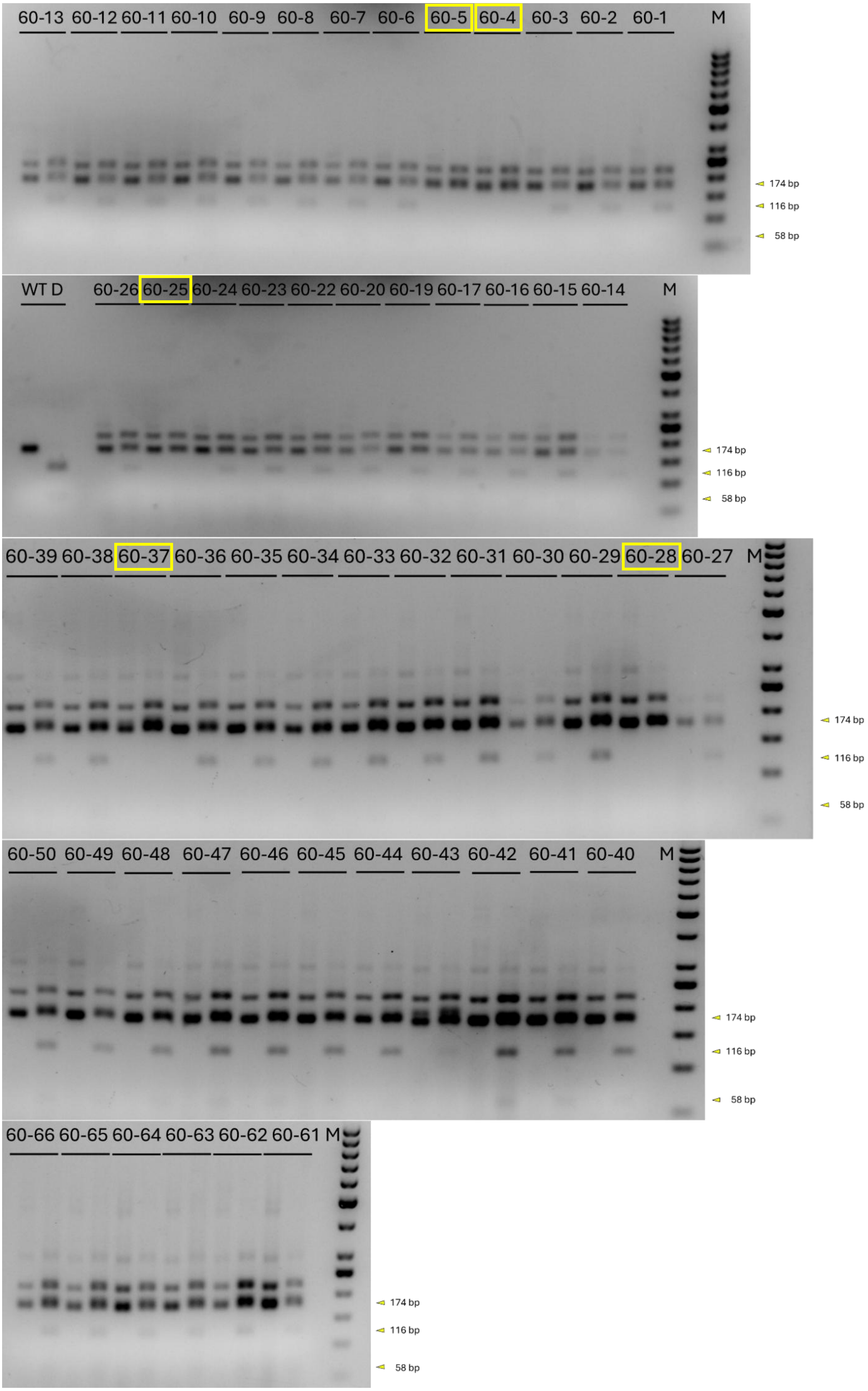

**Fig. S7B: CAPS analysis in cv. Désirée T1 generation - descendants of line 65 (gRNA 1 target site), see the legend below.**

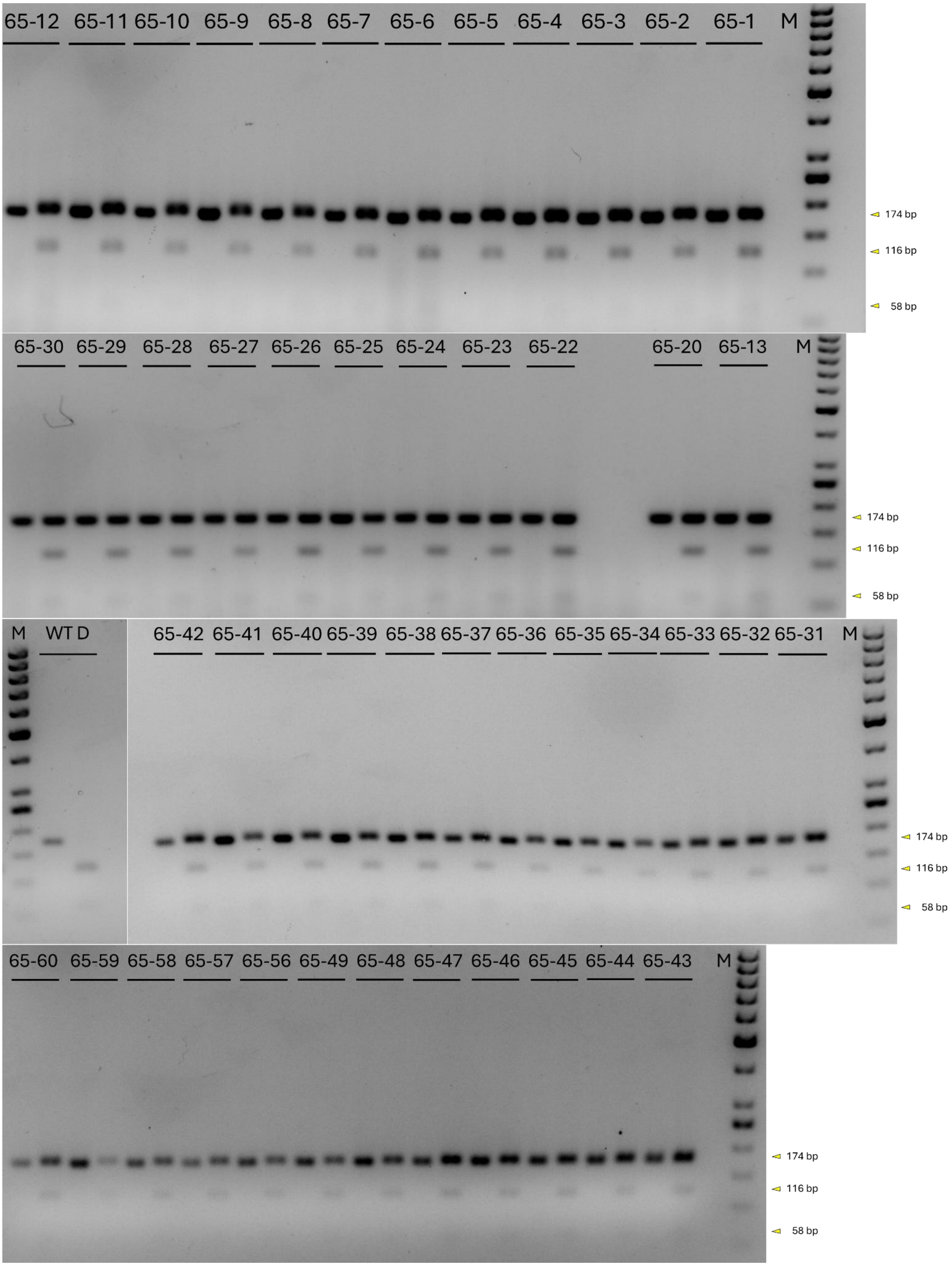

**Fig. S7C: CAPS analysis in cv. Désirée T1 generation - descendants of line 89 (gRNA 1 target site), see the legend below.**

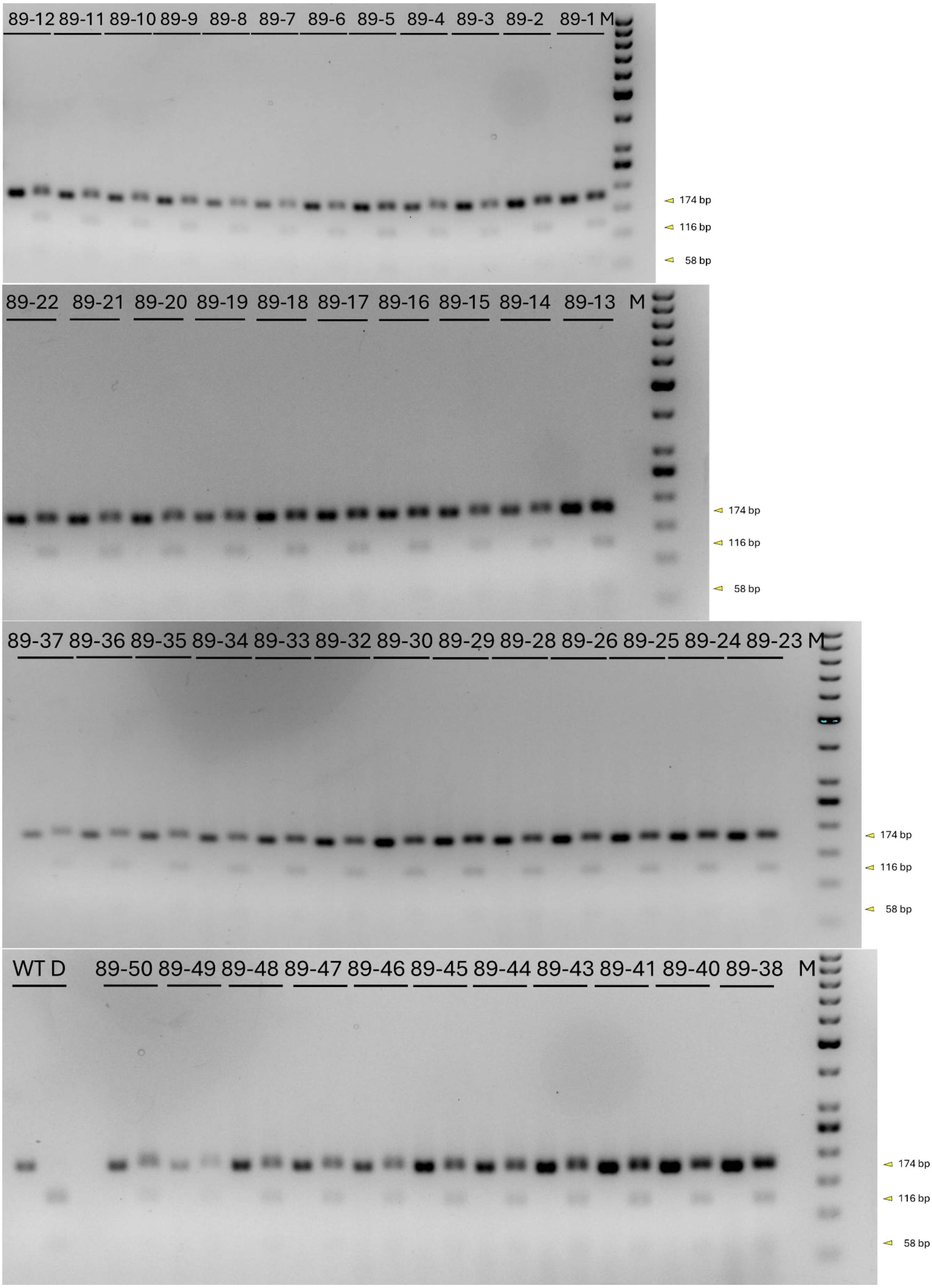

**Fig. S7D: CAPS analysis in cv. Désirée T1 generation - descendants of line 91 (gRNA 1 target site), see the legend below.**

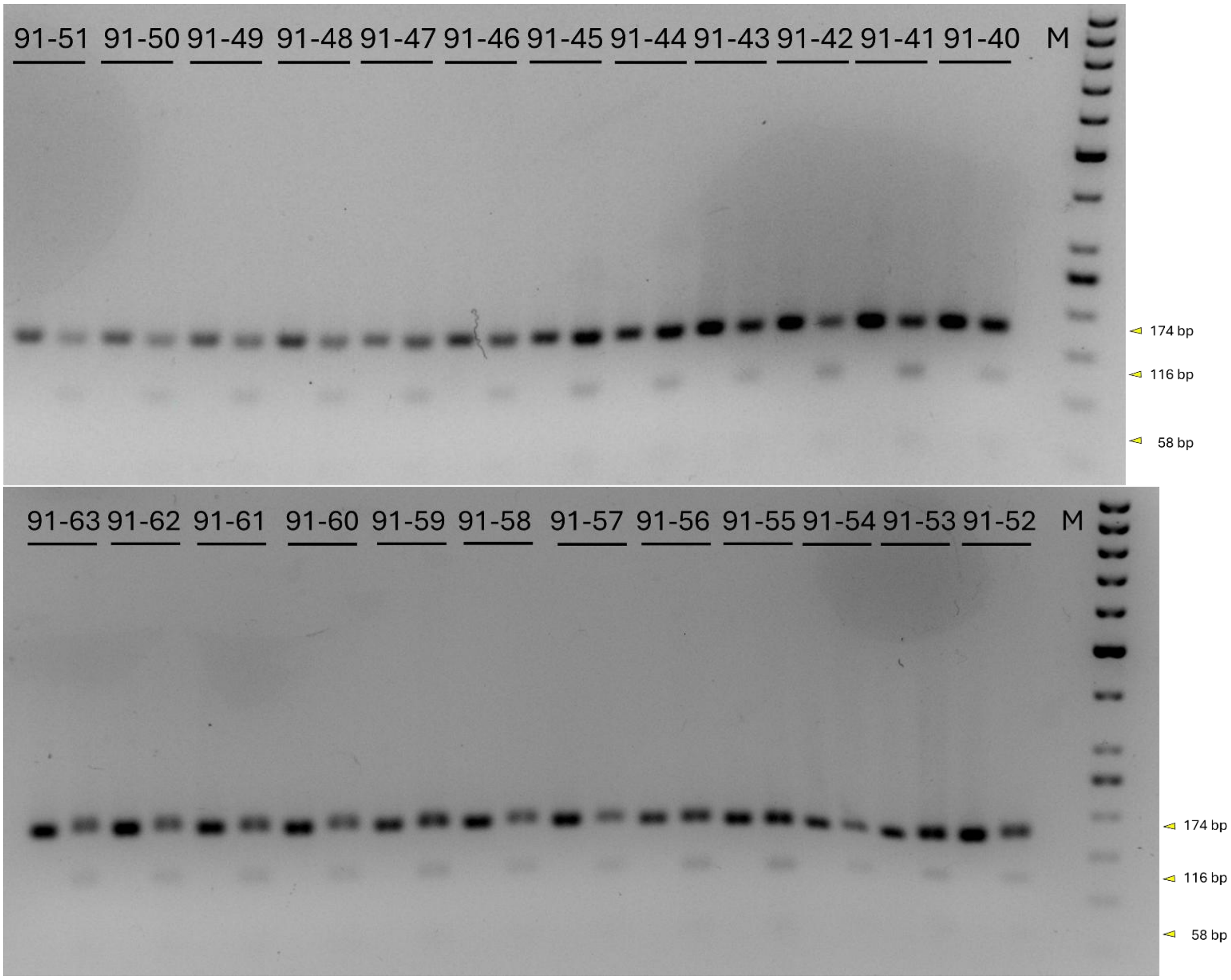

**Fig. S7E: CAPS analysis in cv. Désirée selected lines of T1 generation - (gRNA 2 target site), see the legend below.**

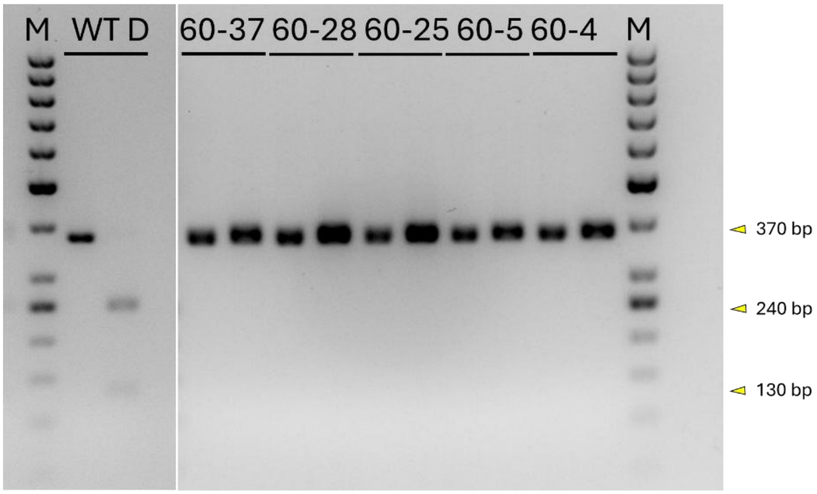


**Fig. S7: CAPS analysis in T1 generation:** lines of cv. Désirée transformed by construct for CRISPR-Cas9 induced knockout of *BEL5* gene – descendants of lines 60 (A), 65 (B), 89 (C), 91 (D) were screened for indels in gRNA1 using SalI. For each transgenic line a pair of samples was tested, from left the first band represents a non-digested control and the second band is a digested sample. WT Désirée (WT D) sample used as positive control showed complete digestion for the SalI site (A) as well as for the HpyF10VI site (B). Most of lines showed both digested and non-digested bands (edited and non-edited DNA), all the descendants of line 60 lines carried the long indel inherited from the progenitor line; five lines (60-4,60-5, 60-25, 60-28 and 60-37, marked with a yellow rectangle) did not show digested products suggesting potential complete editing. Lines (60-4,60-5, 60-25, 60-28 and 60-37) were evaluated also for indels in gRNA 2 site using HpyF10VI (E) and showed just undigested bands suggesting complete editing.

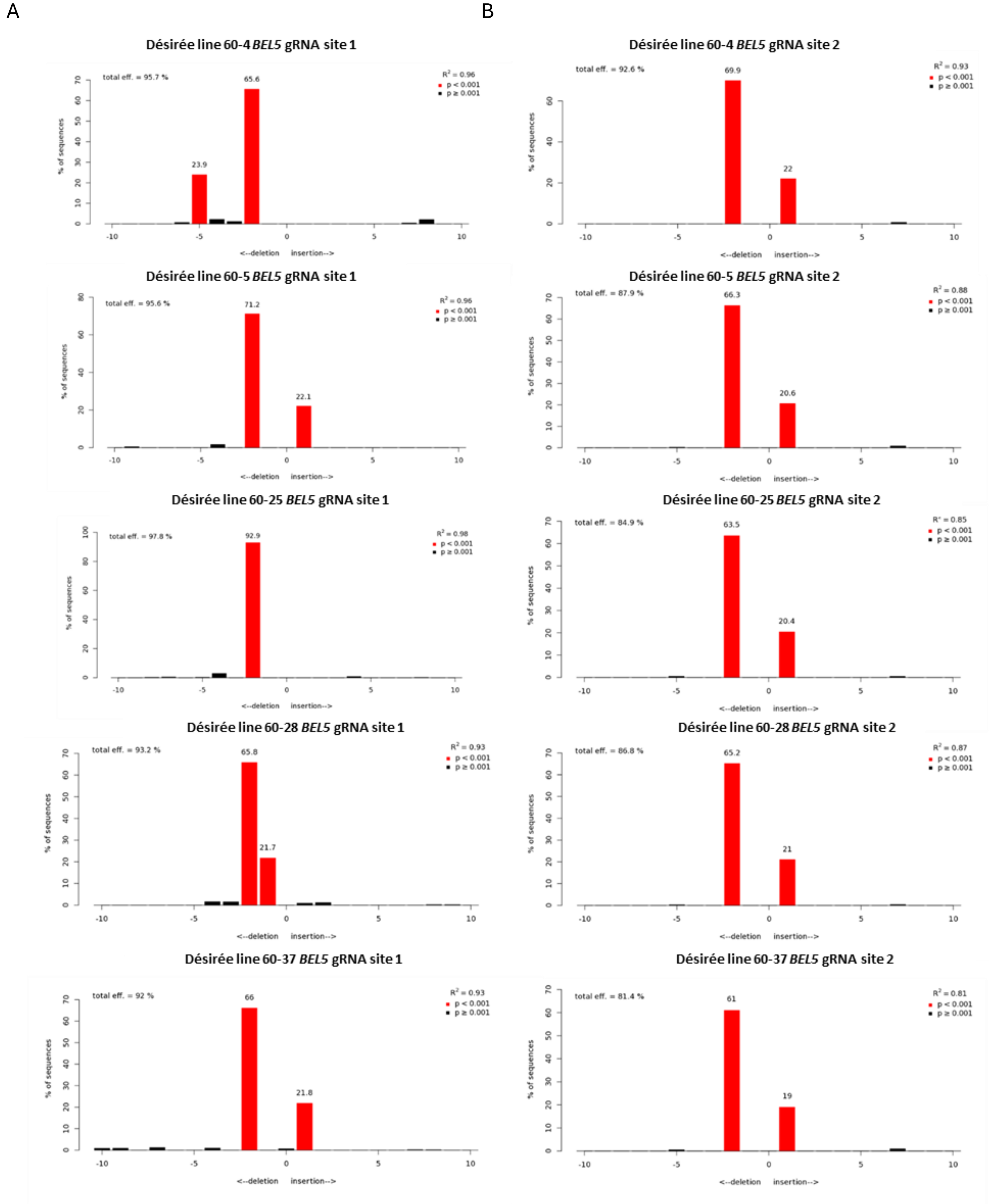
A
B
Désirée line 60-4 *BEL5* gRNA site 1
Désirée line 60-4 *BEL5* gRNA site 2
Désirée line 60-5 *BEL5* gRNA site 1
Désirée line 60-5 *BEL5* gRNA site 2
Désirée line 60-25 *BEL5* gRNA site 1
Désirée line 60-25 *BEL5* gRNA site 2
Désirée line 60-28 *BEL5* gRNA site 1
Désirée line 60-28 *BEL5* gRNA site 2
Désirée line 60-37 *BEL5* gRNA site 1
Désirée line 60-37 *BEL5* gRNA site 2
% of sequences
<--deletion insertion-->

**Fig. S8: Short indels detected of *BEL5* gene in T1 generation lines:** cv. Désirée transformed by construct for CRISPR-Cas9 induced knockout of *BEL5* gene; selected based on CAPS screening (Fig. S7). PCR products covering all alleles were Sanger sequenced and sequence edits and their representation were estimated by TIDE software at gRNA 1 site (A) and gRNA 2 site (B). The obtained results align with the CAPS analysis.

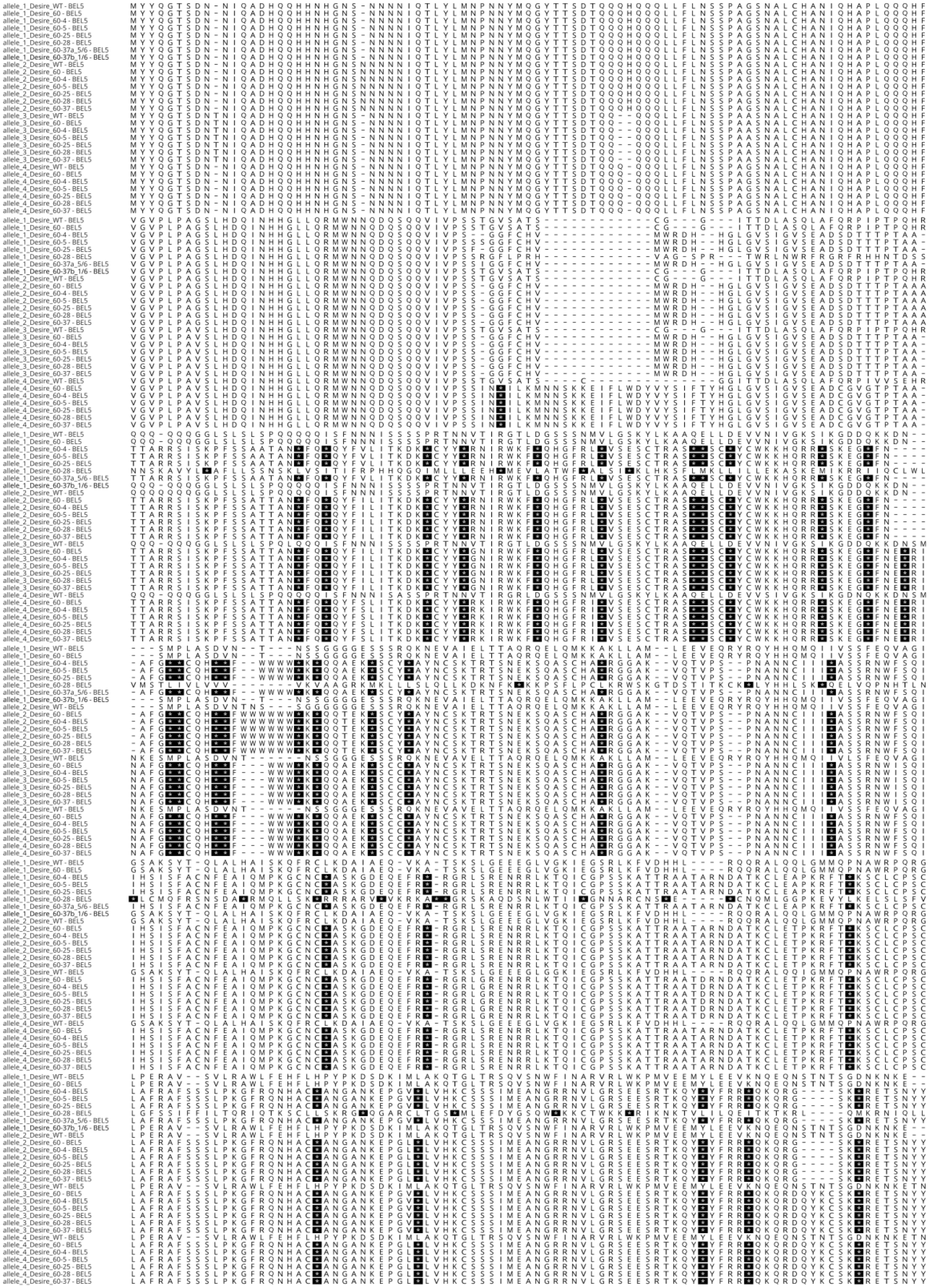

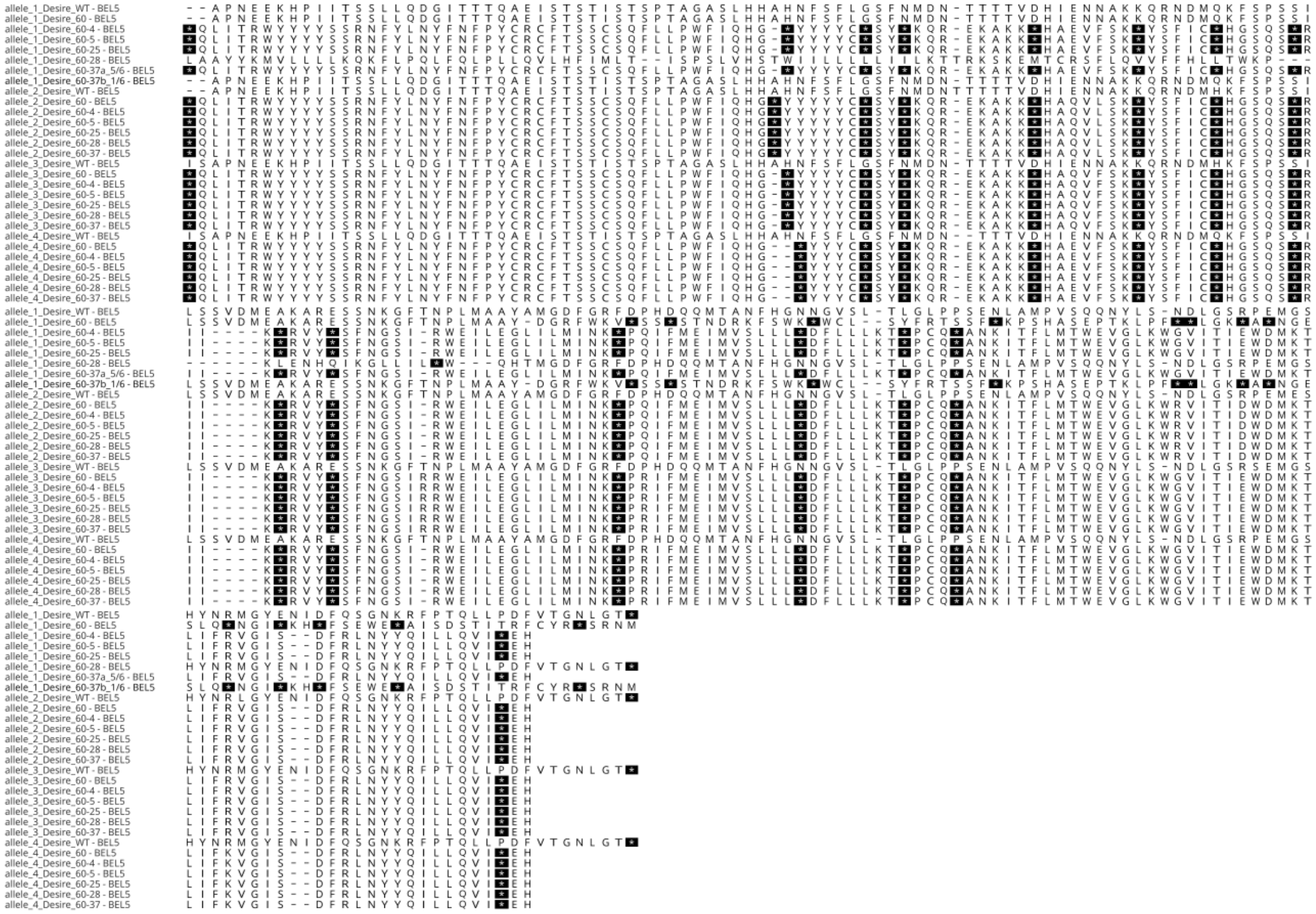

**Fig. S9: Amino acid sequences predicted based on individual *BEL5* alleles:** WT Désirée and lines transformed with the construct for CRISPR–Cas9–induced knockout of the *BEL5* gene — line 60 (T0 generation) and its descendants (T1 generation; lines 60-4, 60-5, 60-25, 60-28, 60-37). Amino acid sequences were predicted from DNA sequences obtained by amplicon sequencing (NGS), which confirmed the mutations predicted by CAPS and Sanger sequencing followed by TIDE analysis, and furthermore revealed a residual portion of unedited sequence at the gRNA1 site in line 60-37, undetectable by the previous methods. Lines 60-4, 60-5, 60-25, and 60-28 carried premature stop codons already in the first exon across all alleles, completely abolishing BEL5 function. While most sequences in line 60–37 also carried stop codons already in the first exon, a small fraction had a stop codon only in the fourth exon, which may not fully disrupt BEL5 function.

**Table S1: List of potato varieties with fully sequenced genomes**

| Genotype/case study | Ploidy | Reference | Genomic data |
|---|---|---|---|
| ***S. tuberosum* group *Phureja* DM1-3 516 R44** | Doubled haploid | Felcher *et al.*, 2012; Hamilton *et al.*, 2011; Pham *et al.*, 2020; The Potato Genome Sequencing Consortium, 2011 | Haplotype-resolved genome assembly |
| **Potato v2.0 project (20 genotypes)** | Doubled haploids | Potato v2.0 project (2020) https://potatov2.github.io/ | Haplotype-resolved genome assemblies |
| ***Solanum chacoense* M6** | Diploid inbred clone | The Buell Lab at the University of Georgia (2022) | Haplotype-resolved genome assembly |
| ***S. tuberosum* group *Tuberosum* RH89-039-16** | Heterozygous diploid | Zhou *et al.*, 2020 | Haplotype-resolved genome assembly |
| ***Solanum candolleanum*** | Diploid | The Buell Lab at the University of Georgia (2022) | Haplotype-resolved genome assembly |
| **Jan** | Diploid | Jiang Lab, Michigan State University (2024) | Haplotype-resolved genome assembly |
| **101 genomes of section *Petota* (including wild species and landraces)**<br>***S. etuberosum* (6 genotypes)** | Diploids | Zhang *et al.*, 2025 | 53- haplotype-resolved genome assemblies<br>3 – haplotype-resolved genome assemblies |
| **67 genotypes (wild species, South American landraces, North American cultivars)** | Diploids and tetraploids | Hardigan *et al.*, 2017 | Illumina reads, not assembled |
| **88 genotypes (landraces, modern cultivars and historical herbarium samples)** | Diploids and tetraploids | Gutaker *et al.*, 2019 | Illumina reads, not assembled |
| **Otava** | Tetraploid | Sun *et al.*, 2022 | Haplotype-resolved genome assembly |
| **Q9 (Qingshu No. 9', 3 875 213 x APHRODITE)** | Tetraploid | Wang *et al.*, 2022 | Haplotype-resolved genome assembly |
| **Cooperation-88 (C88)** | Tetraploid | Bao *et al.*, 2022 | Haplotype-resolved genome assembly |
| **Atlantic, Castle Russet, Avenger, Altus, Colomba, Spunta, Altus x Colomba** | Tetraploids | Hoopes *et al.*, 2022; Serra Mari *et al.*, 2024 | Haplotype-resolved genome assemblies |
| **Diacol Capiro** | Tetraploid | Reyes-Herrera *et al.*, 2024 | Haplotype-resolved genome assembly |
| **Désirée** | Tetraploid | Sevestre *et al.*, 2020; Godec *et al.*, 2025 | Illumina unassembled reads; Haplotype-resolved genome assembly |

Table S2: ***BEL5*** **alleles in** ***S. tuberosum*** **ssp.** ***tuberosum*** **cv. Otava and cv. Désirée**

| **BEL5, cv. Otava** | **J_Browse (https://spuddb.uga.edu/index.shtml), Sun et al., 2022** |
|---|---|
| **Gene ID** | **locus** |
| **He1-St06G504960** | Chr6_He1 Chr6_He1:54582114..54585341 (+ strand) |
| **He2-St06G504760** | Chr6_He2 Chr6_He2:54961363..54967659 (+ strand) |
| **St1-St06G513560** | Chr6_St1 Chr6_St1:54486770..54489817 (+ strand) |
| **St2-St06G489680** | Chr6_St2 Chr6_St2:53416766..53419936 (+ strand) |

| **BEL5, cv. Désirée** | **J_Browse (https://desiree.nib.si), Godec et al., 2025** | |
|---|---|---|
| **this report** | **Gene ID** | **locus** |
| **alelle_1** | **Soltu.Des.v1_C06H3G036870** | chr_06_3:52,323,078..52,327,777 (+) |
| **alelle_2** | **Soltu.Des.v1_C06H4G035220** | chr_06_4:52,546,270..52,551,094 (+) |
| **alelle_3** | **Soltu.Des.v1_C06H1G035050** | chr_06_1:52,781,566..52,786,359 (+) |
| **alelle_4** | **Soltu.Des.v1_C06H2G042900** | chr_06_2:56,832,295..56,837,129 (+) |

**Table S3: Off-target predictions for *BEL5*-targeting gRNAs**. Off-targets in Otava were identified using the CRISPOR tool. For Désirée, off-targets were detected by screening a whole-genome Illumina-reads dataset (SRR8261493) using the Geneious software (see Chapter 4.3); selected reads were subjected to BLAST analysis against the DM reference genome to identify potential off-target genes.

### Otava

| **haplotype** | **gRNA 1** | **gRNA 2** |
|---|---|---|
| | GATAGTACCATCGTCGACGG *GGG* | TTTAATGGCAGCATACGCGA *TGG* |
| He1 | no off-targets in exon | no off-targets in exon |
| He2 | no off-targets in exon | 1 potential off-target in exon: **St06G300640.1**<br>TT**C**AATG**A**CAGC**C**TAC**A**CGA *GGG*<br>CFD Off-target score: 0.084211<br>MIT Off-target score: 0.23 |
| St1 | 1 potential off-target in exon:<br>**St03G356640.1**<br>GA**G**AGT**C**CCATC**A**TCGA**A**GG *TGG*<br>CFD Off-target score: 0.108728<br>MIT Off-target score: 0.08 | no off-targets in exon |
| St2 | no off-targets in exon | 1 potential off-target in exon:<br>**St01G630480.1**<br>TTTAAT**T**GCAG**A**A**C**ACG**A**GA *AGG*<br>CFD Off-target score: 0.062794<br>MIT Off-target score: 0.01 |

### Désirée

| **gRNA1** | **gRNA2** |
|---|---|
| GATAGTACCATCGTCGACGG *GGG* | TTTAATGGCAGCATACGCGA *TGG* |
| potential off targets: | potential off targets: |
| **LOC102597826** | **LOC102599535** |
| G**T**TAGTACC**GAT**GTC**T**ACGG *CGG* | **G**TTAATGG**G**AGCATA**TTT**GA *TGG* |

**LOC102586497**

GAT**G**GT**T**CCATCGT**T**GA**G**G**A** *TGG*

**LOC107058141**

GATAGTACCA**C**CG**C**C**A**A**AT**G *AGG*

**LOC102588882**

GA**C**A**AG**ACCA**C**CGT**G**GACGG *TGG*

**LOC102602026**

**CGGC**GTACCATCGTCG**G**CGG *CGG*

**LOC102595299**

**G**TT**G**AT**T**GCAGCA**A**ACGC**A**A *AGG*

**LOC102585982**

T**A**TAAT**C**GCA**T**CATAC**T**CG**C** *CGG*

**LOC102589990**

T**A**TAATG**AG**A**A**CA**C**ACGCGA *GGG*

**LOC102598882**

T**A**TAATGG**A**AG**A**A**G**A**G**GCGA *GGG*

**LOC102581828**

T**G**TAA**C**GGCAGCA**G**A**A**G**G**GA *GGG*

**LOC102592072**

TT**A**AATGG**TT**GCATAC**CA**GA *TGG*

**LOC102581498**

TTTA**G**T**T**GCAG**G**AT**T**CGC**A**A *GGG*

**LOC102591469**

TTTA**TG**TG**C**AGCATACG**AT**A *CGG*

**Table S4: Commonly used methods for detecting CRISPR-Cas9 mutations in polyploid/mosaic organisms**

| Method | Description and Suitability | Relevant Publications |
|---|---|---|
| **Phenotypic analysis** | Suitable for genes whose knockout results in an obvious phenotype (e.g., albino phenotype from a PDS gene knockout). | Bánfalvi *et al.*, 2020 |
| **Reporter markers** | e.g., GUS or GFP. Primarily used to track transgene expression but can also be inserted into the target locus to indicate editing of a specific allele. Useful for screening but does not reveal exact sequence edits. | Butler *et al.*, 2016; Nakayasu *et al.*, 2018 |
| **Detection of large deletions by PCR** | Large deletions can be easily detected by PCR using primers flanking or specific to the target region. Excision of a segment between two gRNAs may generate such deletions, but it is not very frequent. | Kieu *et al.*, 2021 |

| | | |
|---|---|---|
| **Cleaved amplified polymorphic sequences (CAPS)** | Sensitive even for SNPs, enables estimation of edited vs. non-edited DNA ratio. Requires a restriction site at the target locus. Useful as a screening method but does not provide information about the exact sequence edits in target gene. | Butler *et al.*, 2015; Hegde *et al.*, 2021 |
| **Enzyme mismatch cleavage assays** | Detects heteroduplexes containing mismatches. Typically employs T7 Endonuclease I (T7EI) or Surveyor nuclease (T7EI is generally more sensitive). May not detect mismatches in certain sequence contexts. Useful as a screening method but does not provide information about the exact sequence edits in target gene. | Yang *et al.*, 2017; Zhang *et al.*, 2019 |
| **PCR/RNP method** | *In vitro* digestion of the PCR product with the RNP complex. Useful as a screening method but does not provide information about the exact sequence edits in target gene. | Liang *et al.*, 2018 |
| **High-Resolution Melting Analysis** | Detects mutations by differences in DNA melting curves using fluorescent dyes. Fast and sensitive method useful for screening but does not provide information about the exact sequence edits in target gene. | Lukan *et al.*, 2022 |
| **Sanger sequencing and software analysis** | Chromatograms from Sanger sequencing containing overlapping peaks can be analysed by software (e.g. TIDE or ICE) to decompose the sequence and determine the type and frequency of mutations. | Hegde *et al.*, 2021; Sevestre *et al.*, 2020 |
| **Plasmid cloning and Sanger sequencing** | Cloning of PCR product into a plasmid. Sanger sequencing of individual colonies allows separation of distinct mutation variants. Low representativeness – analysis of a small number of clones, risk of artifacts. | Abeuova *et al.*, 2023; Ly *et al.*, 2023 |
| **Amplicon sequencing (NGS)** | High-throughput sequencing of target amplicons (Illumina, PacBio, Oxford Nanopore). Highly accurate — capable of detecting low-frequency mutations and distinguishing individual alleles. Ideal for final validation, but expensive and requires bioinformatic analysis. | Decima Oneto *et al.*, 2025; Lebedeva *et al.*, 2022 |
| **Droplet Digital PCR** | PCR is performed independently in each droplet, allowing separation of mutation variants and determination of their frequency. Can detect even rare mutations. Requires probes for specific targets and specialized instruments and reagents; not universal; expensive method. | Peng *et al.*, 2020 |

**Table S5: List of primers used in the study**

**The list of primers used for *BEL5* construct assembly**

PCR reaction 1

| | |
|---|---|
| >oCmYLCV F | TGCTCTTCGCGCTGGCAGACATACTGTCCCAC |
| >CSY_gRNA R | TGGTCTCCGCTGCCATTAAACTGCCTATACGGCAGTGAAC |

PCR Reaction 2

| | |
|---|---|
| >REP_gRNA F | TGGTCTCACAGCATACGCGAGTTTTAGAGCTAGAAATAGC |
| >CSY_gRNA R | TGGTCTCCGATGGTACTATCCTGCCTATACGGCAGTGAAC |

PCR Reaction 3

| | |
|---|---|
| >REP_gRNA F | TGGTCTCACATCGTCGACGGGTTTTAGAGCTAGAAATAGC |
| >CSY_term R | TGCTCTTCTGACCTGCCTATACGGCAGTGAAC |

**Primer pair used for amplification of product used for amplicon sequencing**

| | |
|---|---|
| Désiree BEL5 5´UTR F | TTGTTACTTTCTGTTTGCAGGTACTG |
| Désiree BEL5 3´UTR R | AGCTATCAATACGAGACTTTCTGG |

**Primer pair used for detection of the transgene**

| | |
|---|---|
| CmYLCV promoter F | CTAGAAGTAGTCAAGGCGGC |
| 35S terminator R | GCTCAACACATGAGCGAAAC |

**Primers for detection of CRISPR-Cas9 mutations in BEL5 gene**

| | |
|---|---|
| BEL5 gRNA 1 site product F | ACTTTGTCGGTGTGCCTCTT |
| BEL5 gRNA 1 site product R | ATTGAGACGCCAAGTCCGTG |
| BEL5 gRNA 2 site product F | AACAACGCGAAAAAGCAAAGA |
| BEL5 gRNA 2 site product R | TCGGAAATCGCTTATTCCCAC |

**Primers for detection of *Cas9* transcript**

| | |
|---|---|
| Cas9 F | GAGTCTATCCTCCCTAAGAG |
| Cas9 R | CCACAACGAGAACAGAGTAAG |

**Primers for detection of *UBI* transcript (reference control)**

| | |
|---|---|
| UBI F | CTTCAAATTTCTCTTTCAAGATGCAG |
| UBI R | AGCCTTTGCTGATCCGGGG |

**Table S6: Composition of the culture media used in this study.** Growth regulators and antibiotics were added after autoclaving. In the case of antibiotics added to LB, YEB, and MS media, their concentrations are given in the Methods section.

| **Compound** | **Amount in 1 L of medium** |
|---|---|
| | **LB** |
| Peptone | 10 g |
| Yeast extract | 5 g |
| NaCl | 10 g |
| Agar (only for solid LB) | 15 g |
| | **Liquid YEB recovery** |
| Peptone | 10 g |
| Yeast extract | 1 g |
| Sucrose | 68.4 g |
| NaCl | 5.84 g |
| KCl | 1.86 g |
| $MgCl_2 \cdot 6H_2O$ | 20.3 g |
| $MgSO_4 \cdot 7H_2O$ | 24.6 g |
| | **Liquid YEB** |
| Peptone | 10 g |
| Yeast extract | 1 g |
| Sucrose | 5 g |
| $MgSO_4 \cdot 7H_2O$ | 0.5 g |

**Liquid MS**

| | |
|---|---|
| Full MS medium | According to Murashige and Skoog, 1962 |
| Sucrose | 20 g |

| | Callus induction medium | Shoot induction medium |
|---|---|---|
| Full MS medium | According to Murashige and Skoog, 1962 | According to Murashige and Skoog, 1962 |
| Glucose | 16 g | 16 g |
| Myo-inositol | 0,1 g | 0,1 g |
| $ZnSO_4 . 7 H_2O$ | 2 mg | - |
| Agar | 6 g | 6 g |
| NAA | 5 mg | 0,02 mg |
| BAP | 0,1 mg | - |
| $GA_3$ | - | 0,02 mg |
| Trans-zeatin-riboside | - | 2 mg |
| Claforan (Cefotaxime) | 300 mg | 300 mg |
| Kanamycin (selection marker) | 50 mg | 50 mg |

**Solid MS**

| | |
|---|---|
| Full MS medium | According to Murashige and Skoog, 1962 |
| Sucrose | 30 g |
| Agar | 8 g |

**Supplementary references (related to Suppl. Tables 1, 2, 4 and 6)**

Abeuova, L., Kali, B., Tussipkan, D., Akhmetollayeva, A., Ramankulov, Y., and Manabayeva, S. (2023) *CRISPR/Cas9-mediated multiple guide RNA-targeted mutagenesis in the potato*. *Transgenic Research*, **32**, 383–397.

Bánfalvi, Z., Csákvári, E., Villányi, V., and Kondrák, M. (2020) *Generation of transgene-free PDS mutants in potato by Agrobacterium-mediated transformation*. *BMC Biotechnology*, **20**, 25.

Bao, Z., Li, C., Li, G., Wang, P., Peng, Z., Cheng, L., et al. (2022) *Genome architecture and tetrasomic inheritance of autotetraploid potato*. *Molecular Plant*, **15**, 1211–1226.

Butler, N.M., Atkins, P.A., Voytas, D.F., and Douches, D.S. (2015) *Generation and Inheritance of Targeted Mutations in Potato (Solanum tuberosum L.) Using the CRISPR/Cas System*. *PLOS ONE*, **10**, e0144591.

Butler, N.M., Baltes, N.J., Voytas, D.F., and Douches, D.S. (2016) *Geminivirus-Mediated Genome Editing in Potato (Solanum tuberosum L.) Using Sequence-Specific Nucleases*. *Frontiers in Plant Science*, **7**.

Butler, N.M., Jansky, S.H., and Jiang, J. (2020) *First-generation genome editing in potato using hairy root transformation*. *Plant Biotechnology Journal*, **18**, 2201–2209.

Chincinska, I.A., Miklaszewska, M., and Sołtys-Kalina, D. (2023) *Recent advances and challenges in potato improvement using CRISPR/Cas genome editing*. *Planta*, **257**, 25.

Decima Oneto, C.A., Massa, G.A., Echarte, L., Rey Burusco, M.F., Gonzalez, M.N., Alfonso, C.S., et al. (2025) *CRISPR/Cas9 editing of CBP80 enhances drought tolerance in potato (Solanum tuberosum)*. *Frontiers in Plant Science*, **16**, 1598947.

Felcher, K.J., Coombs, J.J., Massa, A.N., Hansey, C.N., Hamilton, J.P., Veilleux, R.E., et al. (2012) *Integration of Two Diploid Potato Linkage Maps with the Potato Genome Sequence*. *PLoS ONE*, **7**, e36347.

Godec, T., Beier, S., Rodriguez-Granados, N.Y., Sasidharan, R., Abdelhakim, L., Teige, M., et al. (2025) *Haplotype-resolved genome assembly of the tetraploid potato cultivar Desiree*. *Scientific Data*, **12**, 1044.

Gutaker, R.M., Weiß, C.L., Ellis, D., Anglin, N.L., Knapp, S., Luis Fernández-Alonso, J., et al. (2019) *The origins and adaptation of European potatoes reconstructed from historical genomes*. *Nature Ecology & Evolution*, **3**, 1093–1101.

Hamilton, J.P., Hansey, C.N., Whitty, B.R., Stoffel, K., Massa, A.N., Van Deynze, A., et al. (2011) *Single nucleotide polymorphism discovery in elite north american potato germplasm*. *BMC Genomics*, **12**, 302.

Hardigan, M.A., Laimbeer, F.P.E., Newton, L., Crisovan, E., Hamilton, J.P., Vaillancourt, B., et al. (2017) *Genome diversity of tuber-bearing* Solanum *uncovers complex evolutionary history and targets of domestication in the cultivated potato*. *Proceedings of the National Academy of Sciences*, **114**.

Hegde, N., Joshi, S., Soni, N., and Kushalappa, A.C. (2021) *The caffeoyl-CoA O-methyltransferase gene SNP replacement in Russet Burbank potato variety enhances late blight resistance through cell wall reinforcement*. *Plant Cell Reports*, **40**, 237–254.

Hoopes, G., Meng, X., Hamilton, J.P., Achakkagari, S.R., De Alves Freitas Guesdes, F., Bolger, M.E., et al. (2022) *Phased, chromosome-scale genome assemblies of tetraploid potato reveal a complex genome, transcriptome, and predicted proteome landscape underpinning genetic diversity*. *Molecular Plant*, **15**, 520–536.

Johansen, I.E., Liu, Y., Jørgensen, B., Bennett, E.P., Andreasson, E., Nielsen, K.L., et al. (2019) *High efficacy full allelic CRISPR/Cas9 gene editing in tetraploid potato*. *Scientific Reports*, **9**, 17715.

Kieu, N.P., Lenman, M., Wang, E.S., Petersen, B.L., and Andreasson, E. (2021) *Mutations introduced in susceptibility genes through CRISPR/Cas9 genome editing confer increased late blight resistance in potatoes*. *Scientific Reports*, **11**, 4487.

Lebedeva, M., Komakhin, R., Konovalova, L., Ivanova, L., Taranov, V., Monakhova, Y., et al. (2022) *Development of potato (Solanum tuberosum L.) plants with StLEAFY knockout*. *Planta*, **256**, 116.

Lee, S.-Y., Kang, B., Venkatesh, J., Lee, J.-H., Lee, S., Kim, J.-M., et al. (2024) *Development of virus-induced genome editing methods in Solanaceous crops*. *Horticulture Research*, **11**, uhad233.

Liang, Z., Chen, K., Yan, Y., Zhang, Y., and Gao, C. (2018) *Genotyping genome-edited mutations in plants using CRISPR ribonucleoprotein complexes*. *Plant Biotechnology Journal*, **16**, 2053–2062.

Lukan, T., Veillet, F., Križnik, M., Coll, A., Mahkovec Povalej, T., Pogačar, K., et al. (2022) *CRISPR/Cas9-mediated fine-tuning of miRNA expression in tetraploid potato*. *Horticulture Research*, **9**, uhac147.

Ly, D.N.P., Iqbal, S., Fosu-Nyarko, J., Milroy, S., and Jones, M.G.K. (2023) *Multiplex CRISPR-Cas9 Gene-Editing Can Deliver Potato Cultivars with Reduced Browning and Acrylamide*. *Plants*, **12**, 379.

Murashige, T. and Skoog, F. (1962) *A Revised Medium for Rapid Growth and Bio Assays with Tobacco Tissue Cultures*. *Physiologia Plantarum*, **15**, 473–497.

Nakayasu, M., Akiyama, R., Lee, H.J., Osakabe, K., Osakabe, Y., Watanabe, B., et al. (2018) *Generation of α-solanine-free hairy roots of potato by CRISPR/Cas9 mediated genome editing of the St16DOX gene*. *Plant Physiology and Biochemistry*, **131**, 70–77.

Park, J.-S., Park, K.H., Park, S.-J., Ko, S.-R., Moon, K.-B., Koo, H., et al. (2023) *WUSCHEL controls genotype-dependent shoot regeneration capacity in potato*. *Plant Physiology*, **193**, 661–676.

Peng, C., Zheng, M., Ding, L., Chen, X., Wang, X., Feng, X., et al. (2020) *Accurate Detection and Evaluation of the Gene-Editing Frequency in Plants Using Droplet Digital PCR*. *Frontiers in Plant Science*, **11**, 610790.

Pham, G.M., Hamilton, J.P., Wood, J.C., Burke, J.T., Zhao, H., Vaillancourt, B., et al. (2020) *Construction of a chromosome-scale long-read reference genome assembly for potato*. *GigaScience*, **9**, giaa100.

Reyes-Herrera, P.H., Delgadillo-Duran, D.A., Flores-Gonzalez, M., Mueller, L.A., Cristancho, M.A., and Barrero, L.S. (2024) *Chromosome-scale genome assembly and annotation of the tetraploid potato cultivar Diacol Capiro adapted to the Andean region*. *G3: Genes, Genomes, Genetics*, **14**, jkae139.

Serra Mari, R., Schrinner, S., Finkers, R., Ziegler, F.M.R., Arens, P., Schmidt, M.H.-W., et al. (2024) *Haplotype-resolved assembly of a tetraploid potato genome using long reads and low-depth offspring data*. *Genome Biology*, **25**, 26.

Sevestre, F., Facon, M., Wattebled, F., and Szydlowski, N. (2020) *Facilitating gene editing in potato: a Single-Nucleotide Polymorphism (SNP) map of the Solanum tuberosum L. cv. Desiree genome*. *Scientific Reports*, **10**, 2045.

Sun, H., Jiao, W.-B., Krause, K., Campoy, J.A., Goel, M., Folz-Donahue, K., et al. (2022) *Chromosome-scale and haplotype-resolved genome assembly of a tetraploid potato cultivar*. *Nature Genetics*, **54**, 342–348.

The Potato Genome Sequencing Consortium (2011) *Genome sequence and analysis of the tuber crop potato*. *Nature*, **475**, 189–195.

Vinterhalter, D., Zdravković-Korać, S., Banjac, N., Dragićević, I.Č., Cingel, A., Raspor, M., and Ninkovi, S. (2008) *Protocols for Agrobacterium-mediated Transformation of Potato*. *Fruit, Vegetable and Cereal Science and Biotechnology*, 1–15.

Wang, F., Xia, Z., Zou, M., Zhao, L., Jiang, S., Zhou, Y., et al. (2022) *The autotetraploid potato genome provides insights into highly heterozygous species*. *Plant Biotechnology Journal*, **20**, 1996–2005.

Yang, H., Wu, J.-J., Tang, T., Liu, K.-D., and Dai, C. (2017) *CRISPR/Cas9-mediated genome editing efficiently creates specific mutations at multiple loci using one sgRNA in Brassica napus*. *Scientific Reports*, **7**, 7489.

Zhang, Z., Hua, L., Gupta, A., Tricoli, D., Edwards, K.J., Yang, B., and Li, W. (2019) *Development of an* Agrobacterium *-delivered CRISPR /Cas9 system for wheat genome editing*. *Plant Biotechnology Journal*, **17**, 1623–1635.

Zhang, Z., Zhang, P., Ding, Y., Wang, Z., Ma, Z., Gagnon, E., et al. (2025) *Ancient hybridization underlies tuberization and radiation of the potato lineage*. *Cell*, **188**, 5249-5265.e15.

Zhou, Q., Tang, D., Huang, W., Yang, Z., Zhang, Y., Hamilton, J.P., et al. (2020) *Haplotype-resolved genome analyses of a heterozygous diploid potato*. *Nature Genetics*, **52**, 1018–1023.
